# Spatiotemporal Decomposition of Whole-Brain Alpha Traveling Waves

**DOI:** 10.1101/2024.08.23.609472

**Authors:** Yichao Li, Bo Hong

## Abstract

Spontaneously emerging traveling waves are present within the spatiotemporal patterns of alpha-band EEG oscillations, but current analysis methods are limited in parsing the diversity of global wave structures and their correlation with brain functions. To address this limitation, we constructed a rigorous mathematical framework, Weakly Orthogonal Conjugate Contrast Analysis (WOCCA), which decomposes the whole-brain EEG alpha oscillations into directionally independent traveling waves. For the first time, we systematically characterized propagating components in alpha-band resting-state EEG as a combination of rotational, longitudinal, and horizontal traveling wave patterns. The intensity, directionality, and morphological characteristics of these wave patterns account for the differences between cognitive states during rest and consciousness levels under sedation. Moreover, our WOCCA decomposition encompassed the state transition dynamics captured by EEG Microstate Analysis, a conventional analysis framework for alpha waves. These results not only established a novel approach for identifying and analyzing traveling waves but also provided evidence for the relationship between wave directionality and cooperative interactions in brain network.

## Introduction

Ever since Hans Berger first recorded brain signals in 1924, alpha waves have been a ubiquitous phenomenon in electroencephalograph (EEG) recordings (**Berger, 1929**). Characterized by prominent oscillations within the 7-13 Hz frequency range, alpha wave amplitudes increase and decrease with eye closure and opening (**Adrian and Matthews, 1934**). The widespread presence of alpha waves and their significant relation to cognitive functions have rendered them a crucial research object in neuroscience for nearly a century. The spatiotemporal structure of alpha waves, including the spatial distribution of amplitude (**Jokisch and Jensen, 2007**; **Haegens et al., J Neurosci., 2011**) and instantaneous local phase (**Mathewson et al., 2009**; **Hansen et al., 2019**), have been found to influence human attention and decision-making behaviors. The underlying mechanism is believed to be the local neural population excitability modulated by oscillating alpha wave potentials (**Schroeder and Lakatos, 2009**; **Jensen and Mazaheri, 2010**).

EEG Microstate Analysis (**Pascual-Marqui et al., 1995**) is a popular framework for describing spatiotemporal structure of alpha waves. The instantaneous spatial topography of alpha waves can be abstracted as a series of microstate labels that are discrete in both space and time (**Lehmann et al., 1987**; **Michel and Koenig, 2017**) (**Figure 1a**). The time series of these microstate labels have been widely used to establish neural markers for cognitive functions (**Bréchet et al., 2019**; **Zanesco et al., 2020**) and pathological states (**da Cruz et al., 2020**; **Gui et al., 2020**). However, the mechanisms underlying generation and physiological functions of microstates lacks a concrete connection with the mainstream theory of neural population modulation by alpha wave. The microstate analysis methodology is also criticized for discreteness, which is incompatible with the continuous nature of alpha oscillations (**Mishra et al., 2020**).

**Figure 1:**
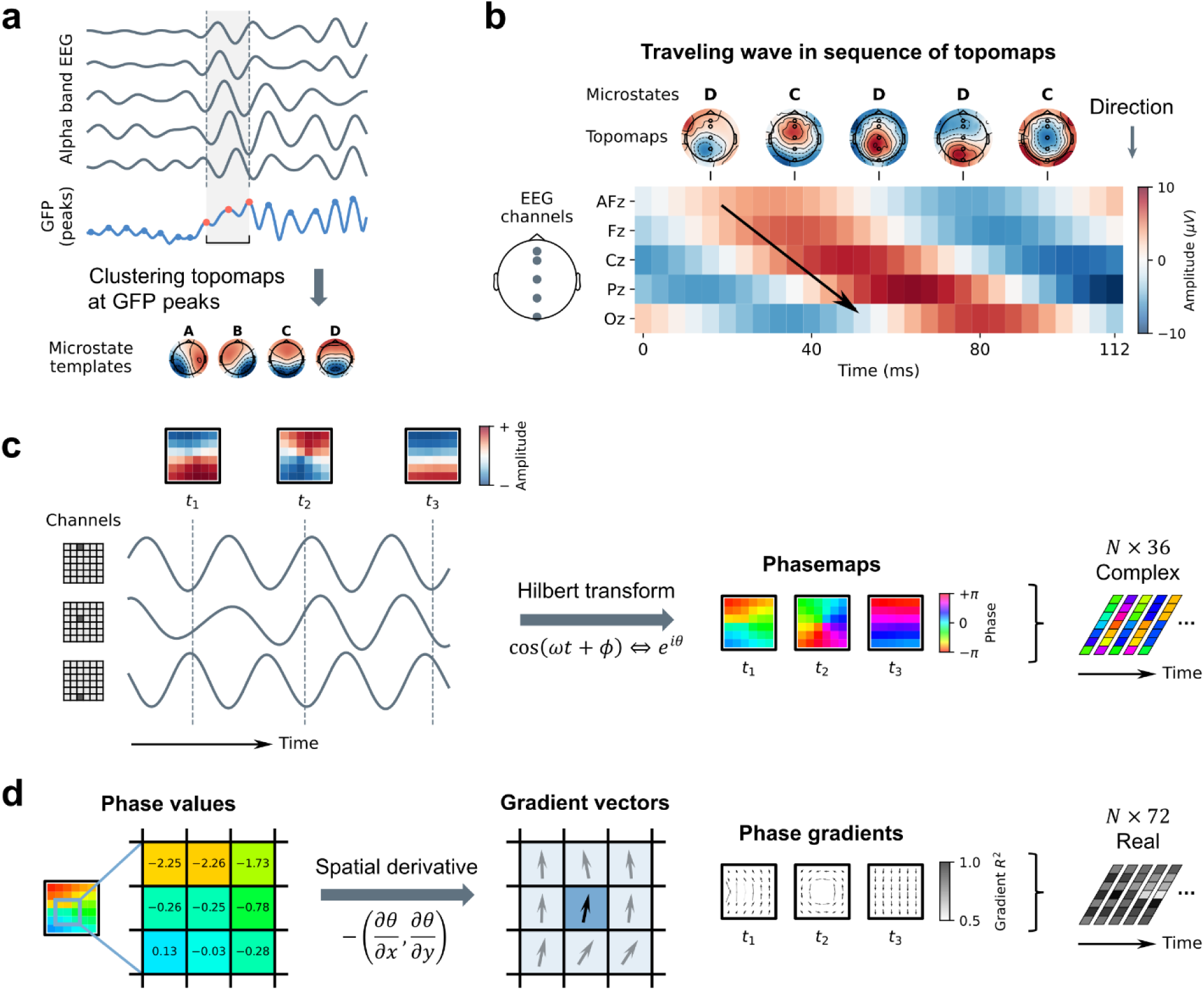
Parsing Traveling Waves in Neural Oscillations. a. Standard procedure of microstate analysis of alpha-band EEG. Microstate templates are chosen as cluster centers of EEG topographies at Global Field Power (GFP) peaks. b. In a short segment of alpha-band EEG (gray shaded area in **a**), transitions between microstate labels C and D (top) represents movement of electric potential peaks from anterior to posterior electrodes, corresponding to a backward longitudinal traveling wave. c. Using synthetic data on a 6×6 grid as an example. Multi-channel oscillation data can be transformed to phasemaps represented by complex-valued vectors using Hilbert transform, which depict relative phase relationships between different channels in a traveling wave. d. Computing the spatial gradient of the phasemaps yields the phase gradient vector fields that directly characterize the direction of wave propagation. Vector grayscale encodes and local coherence of phase gradients (measured by *R*^2^).

The self-emergent traveling wave phenomenon in alpha waves (**Figure 1b**) may provide a more natural perspective. In a short oscillatory segment, a movement in the electric potential topography is essentially a spatial pattern of phase shift (**Nunez, 1974**), making the whole-brain instantaneous phase distribution, or "phasemap" (**Figure 1c**), an ideal candidate for the basic unit composing alpha traveling waves. On the other hand, the "phase gradient field" (**Townsend and Gong, 2018**) visualizes the directionality of wave propagation at each spatial location in a straightforward way (**Figure 1d**). Under the theory of neural population modulation by alpha waves, propagation and directionality, being unique to traveling waves, are deeply connected to the spatiotemporal modes of multi-area neural population activation. This mechanism has been widely used to explain inter-areal brain communication (**Aggarwal et al., 2022**; **Liang et al., 2023**) as well as neural population excitability and cooperation (**Davis et al., 2020**; **Dickey et al., 2021**).

Most existing studies on Alpha traveling waves in scalp EEG focus on the prominent phenomenon of forward and backward propagation, using methods such as phase difference analysis (**Sauseng et al., 2005**; **Fellinger et al., 2012**), phase gradient analysis (**Patten et al., 2012**), or 2D-FFT combining space and time domain (**Alamia and VanRullen, 2019**; **Alamia et al., 2023**) to identify wave directionality. The results generally show that the asymmetry of forward and backward propagation directionality is related to cognitive states and behavioral performance, with possible connections to information processing in bottom-up and top-down directions, respectively (**Klimesch et al., 2006**), suggesting that wave directionality may be an important mechanism for directed neural computation pathways. Nonetheless, considering high complexity of information processing in human brain, traveling waves related to computational functions are likely to exhibit a similar level of diversity. Therefore, analyzing only a pair of pre-assumed wave directions doesn’t guarantee a comprehensive understanding of the global structure of alpha traveling waves, nor does it facilitate further validation of relationships between multiple wave directions and cognitive functions.

Analyzing multi-channel oscillatory data for multiple global wave modes proves to be a challenging task itself, and there are currently no such studies on scalp EEG. Existing methods designed for other neuroimaging modalities, such as Complex Principal Component Analysis (CPCA) on phasemaps (**Bolt et al., 2022**) or Vector Field Singular Value Decomposition (VSVD) on phase gradient fields (**Townsend and Gong, 2018**), use classic matrix decomposition methods to extract the most representative wave components, but they are limited in either parsing wave directionality or separating non-orthogonal propagation modes by their fundamental mathematical properties. To address these limitations, we propose a novel framework named "Weakly Orthogonal Conjugate Contrast Analysis" (WOCCA) for extracting dominant traveling wave patterns from multi-channel EEG data, aiming at maximizing difference in wave directionality. As a result, our method combines the strengths of CPCA and VSVD while also has more general applicability (**Figure 2**). Using WOCCA, we validated the traveling nature of alpha waves and robustly obtained dominant traveling wave patterns with diverse spatiotemporal organization, including anterior-posterior (longitudinal), left-right (horizontal), and rotational waves (**Figure 3**).

**Figure 2:**
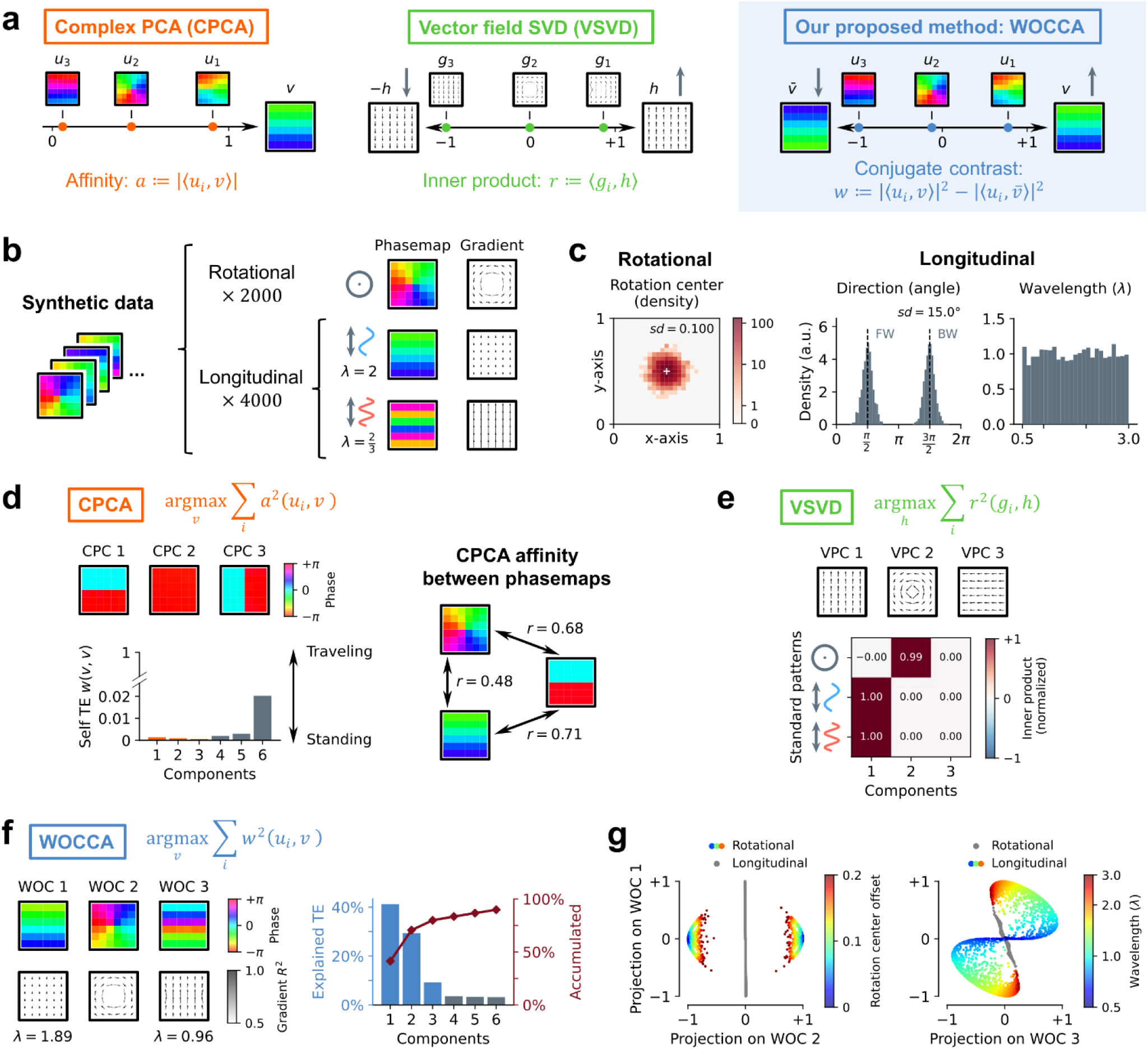
Identifying Traveling Waves Using WOCCA. a. Schematic of methods for measuring and identifying traveling waves. Left: Complex Principal Component Analysis (CPCA) on phasemaps. Middle: Vector Field Singular Value Decomposition (VSVD) on phase gradient vector fields. Right: Our proposed method, Weakly Orthogonal Conjugate Contrast Analysis (WOCCA), using the conjugate contrast function of phasemaps as a metric of similarity between traveling waves. b. Synthetic dataset designed to test the three measure and identification methods, comprising rotational waves as well as longitudinal waves of varying wavelengths, with random perturbations. c. Statistical properties of the synthetic dataset. Left: distribution of rotation center positions of rotational waves; Right: distribution of propagation directions and wavelengths (with the side length square canvas scaled to 1) of longitudinal waves. d. Upper left: the first three principal components extracted by CPCA from phasemaps in the synthetic dataset. Lower left: these principal components exhibit very little traveling wave characteristic measured by self-conjugate contrast function (i.e. self-traveling energy), similar to standing waves. Right: a rotational traveling wave, a longitudinal traveling wave, and a standing wave show high similarity measure by inner product, thus inseparable from each other under orthogonality constraint. e. Upper: the first three principal components extracted by VSVD from phase gradient fields in the synthetic dataset. Lower: longitudinal waves of different wavelengths are indistinguishable under the normalized inner product metric. f. Left: the first three weakly orthogonal components (WOCs) extracted by WOCCA from phasemaps in the synthetic dataset, with Components 1 and 3 corresponding to long- and short-wavelength longitudinal traveling waves, respectively. Right: proportion of total traveling energy in the dataset explained by each component. g. WOCs reveal fine structures of traveling waves: Projections on Component 2 separate rotational waves from longitudinal waves in the dataset, with its magnitude indicating the distance of the rotation centers from the canvas center (left); The plane formed by Components 1 and 3 projections separate longitudinal waves of different wavelengths (right).

As a potential perspective for understanding alpha wave physiology and computational functions, the spatial and temporal continuity of traveling waves has been found to be useful in describing formation of microstate spatial patterns (**Gabay et al., 2018**) as well as their repetitive segments and autocorrelation (**Von Wegner et al., 2021**). However, the rich physiological features contained in microstate sequence dynamics have not yet been explored from the perspective of traveling waves. Our study, based on WOCCA-decomposed alpha traveling wave patterns in resting state and sedated state EEG data, discovered systematic differences between dominant traveling waves in two basic cognitive states of eyes-closed and eyes-open (**Figure 4**) and defined traveling wave markers for gradually decreasing consciousness levels (**Figure 5**), thereby establishing a strong connection between traveling wave patterns and directionality with brain communication and information flow under different cognitive states. Compared with discrete snapshots captured by classic microstate analysis, our method treats EEG data as continuous segments or “clips” of traveling waves. By assigning temporally discrete microstate labels to the dynamics traveling waves, we demonstrated that microstate transition dynamics can be explained by directionality of the ongoing traveling wave (**Figure 6**). This provides a new approach for interpreting the relationship between microstate statistics and brain function.

## Results

### The WOCCA algorithm identifying traveling wave patterns

To evaluate different methods’ ability to identify dominant traveling wave patterns in a relatively realistic scenario, we synthesized traveling wave phasemaps on a virtual 6×6 grid, along with their corresponding phase gradient fields (**Fig. 2b**). The phasemap dataset comprises rotational waves around a central singularity and longitudinal waves in forward or backward directions, with rotation center positions and the angles of propagation direction randomly perturbed. Additionally, the longitudinal waves have randomly distributed wavelengths (**Fig. 2c**). We expect an ideal traveling wave decomposition method to be able to identify rotational and longitudinal wave structures from the dataset and recover important parameters such as wavelength.

We first evaluated the existing CPCA (**Bolt et al., 2022**) and VSVD (**Townsend and Gong, 2018**) methods. For a given set of phasemaps {*u*_*i*_}, the CPCA algorithm obtains the Complex Principal Components (CPCs) *v*_*k*_ by maximizing the sum of squared amplitude of its inner product with the dataset, and constrains the components to be pairwise orthogonal:

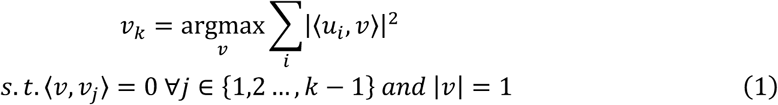

On the synthetic dataset, the first three principal components of CPCA exhibited more characteristics of standing waves than traveling waves (**Fig. 2d left**). An obvious reason is that inner product and orthogonality are (counterintuitively) unsuitable for measuring distance between complex-valued phasemaps. Note that rotational and longitudinal phasemaps are non-orthogonal, and both have high similarity with anti-phase standing waves (**Fig. 2d right**). This makes it impossible to identify these two types of traveling waves from the data using PCA-based phasemap decomposition methods, and causes generation of spurious standing wave components (see **Supplementary Note 4** for a more detailed explanation).

For a given set of phase gradient fields {*g*_*i*_}, the VSVD algorithm obtains the principal components ℎ_*k*_of vector fields by maximizing the sum of squared its normalized inner products with the dataset, also subject to orthogonality constraints:

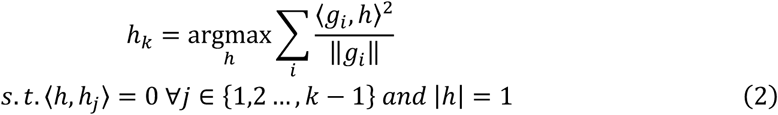

The normalized inner product ranges from −1 to 1, thus holding directionality information for distinguishing traveling waves in the same or opposite directions. On the synthetic dataset, the first two Vector Field Principal Components (VPCs) correctly identified a rotational and a longitudinal wave pattern. However, due to natural limitation of the gradient field representation of traveling waves, it is unable to distinguish longitudinal waves with different wavelengths (**Fig. 2e**, also see **Supplementary Figure 1**). Moreover, the gradient field representation in general cannot be converted back to phasemap form, hence cannot be used for interpreting spatial topography information (such as microstates) in traveling waves.

To circumvent limitations of the two methods above, we propose a novel framework named "Weakly Orthogonal Conjugate Contrast Analysis" (WOCCA). The motivation is that under phasemap representation of traveling waves, the operation of complex conjugation on phasemap vectors is effectively reversing all phase differences between channels. This is naturally convenient for representing wave directionality, since applying complex conjugation to a phasemap is equivalent to flipping wave direction. Therefore, we construct the conjugate contrast function *w*(*u*, *v*) as the squared amplitude of inner product between a phasemap of interest and a target phasemap, minus the inner product between itself and the target’s complex conjugate. Similar to VSVD, this method maps the phasemap of interest to the range of −1 to 1, effectively representing its relative position between the target phasemap’s direction and the opposite one:

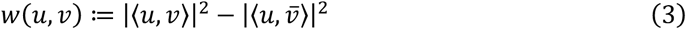

WOCCA obtains the dominant Weakly Orthogonal Components (WOCs) by maximizing the sum of squared conjugate contrast between the components and the dataset. The WOCs are subject to weak orthogonality constraints, i.e., the conjugate contrasts between WOCs are all zero:

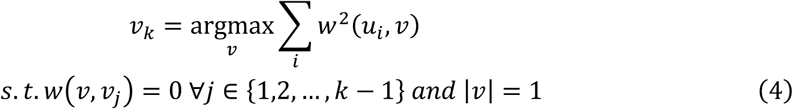

We have rigorously proved that WOCCA possesses two critical properties. First, similar to PCA’s independent decomposition of variance, WOCCA essentially decomposes the "traveling energy" in the dataset, defined as the sum of squared conjugate contrasts of each phasemap with itself, into independent components. Second, for two oscillation segments represented by two weakly orthogonal phasemaps, any one of them contains at least one frame (i.e., an instantaneous topography) that is orthogonal to all frames in the other segment, thereby ensuring distinctiveness of two propagation modes. Mathematical details and proofs are provided in **Supplementary Notes 1-3**. On the synthetic dataset, WOCCA’s first three dominant WOCs consist of one rotational wave and two longitudinal waves with different wavelengths, accounting for 79.8% of total traveling energy (**Fig. 2f**). Moreover, by projecting the data onto the axes of the three components, the separation of wave patterns with different rotation center positions and wavelengths can be clearly visualized (**Fig. 2g**), thus the fine structures within the synthetic dataset are successfully resolved by WOCCA.

### Alpha traveling waves at resting state

The focus of our study is alpha oscillations at resting state. We applied WOCCA to two different EEG datasets: Dataset 1 (**Babayan et al., 2019**) contains alternating epochs of Eyes-Closed (EC) and Eyes-Open (EO) resting state, while Dataset 2 (**Wang et al., 2022**) consists of continuous epochs of EC and EO, recorded in three separate sessions. Although their electrode layouts are different, the WOCs obtained from the two datasets are similar: the first component is rotational around a singularity close to Cz channel; the second component propagates in longitudinal direction along the anterior-posterior central axis; the third component propagates horizontally along the left-right C3-Cz-C4 axis; and the fourth component propagates laterally, also in forward-backward direction but along the left and right hemispheres rather than the central axis (**Fig. 3b, 3d**). The proportion of traveling energy explained by the first four WOCs of the two datasets are also similar, dominated by rotational and longitudinal components. From the fifth component onward, although their contribution to explained traveling energy didn’t exhibit a steep drop, their phasemaps lose whole brain-level coherency (e.g., both the anterior half and the posterior half of the fifth component exhibits horizontal phase gradient, but between them there is a boundary of discontinuity with 180° phase reversal. Also see **Supplementary Figure 2**). Therefore, our subsequent analysis will focus on the first four WOCs.

**Figure 3:**
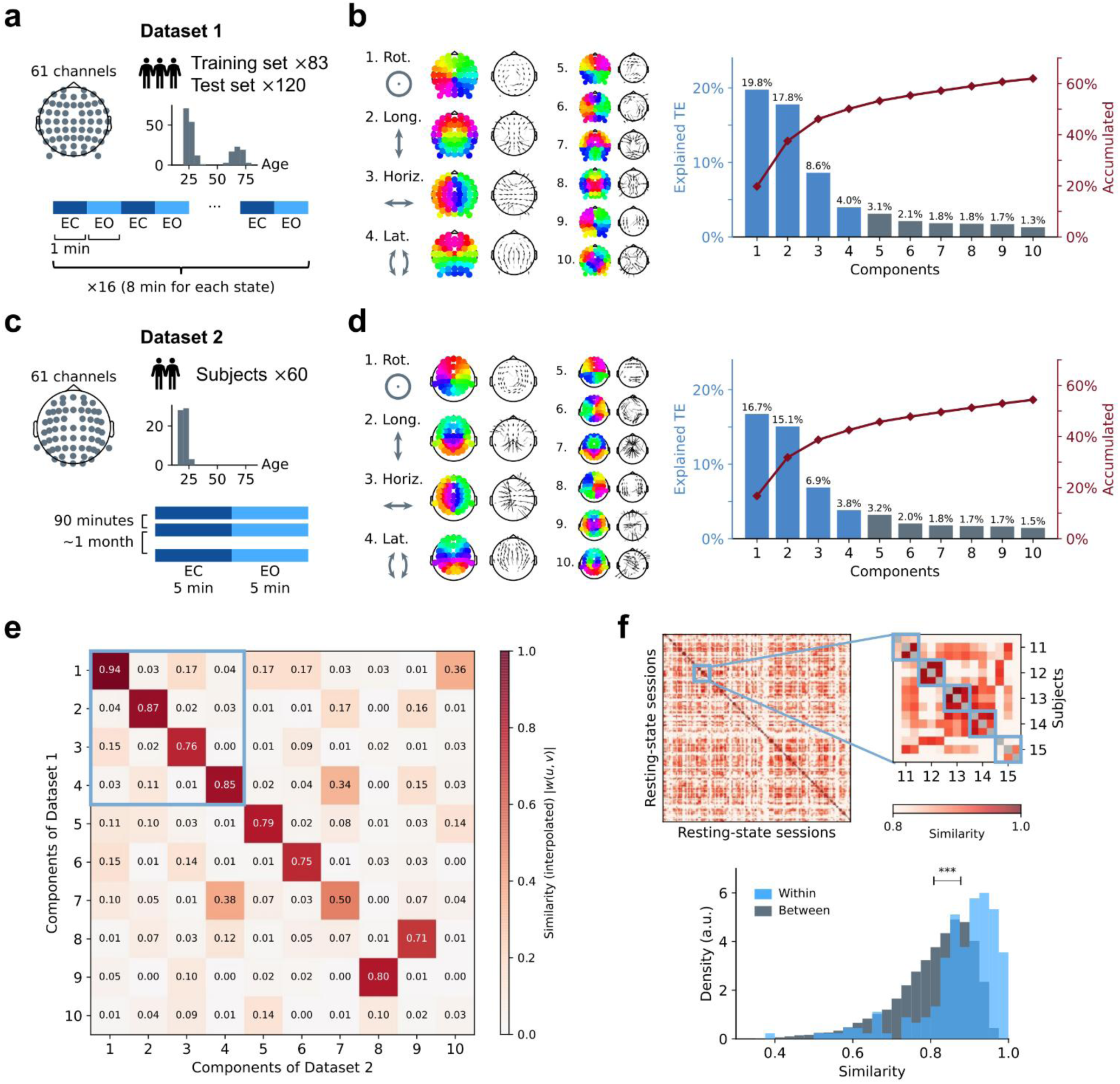
Dominant Traveling Wave Patterns in Resting State EEG. a. Overview of the Mind-Brain-Body Dataset (**Babayan et al., 2019**; referred to as Dataset 1). During resting state EEG recordings, each subject alternately performed 8 rounds of 1-minute Eyes-Closed (EC) and Eyes-Open (EO) rest. Subjects with no bad channels in EEG recording were assigned to the training split, while the remaining subjects comprised the test split. b. Left: WOCs obtained by WOCCA from training split of Dataset 1, with the first four corresponding to rotating, longitudinal, horizontal, and lateral patterns. Right: proportion of total traveling energy in the dataset explained by each component. c. Description of the Test-Retest Resting and Cognitive State EEG Dataset (**Wang et al., 2022**; referred to as Dataset 2). In resting state EEG recordings, each subject underwent 5 minutes of EC and 5 minutes of EO rest per session, with second and third sessions conducted 90 minutes after the first session and approximately one month later, respectively. Although both datasets contain 61 EEG channels, their electrode layouts are different. d. Analogous to **b**, but for analysis of Dataset 2. e. Similarity between WOCs across the two datasets (via spatial interpolation to bridge the differing electrode layouts), measured by the absolute value of the conjugate contrast. f. Within-subject stability of WOCs, with WOCCA applied separately to EEG data of each session of each subject in Dataset 2. Top: overall similarity of the first four WOCs across sessions and subjects, with each 3×3 diagonal block (excluding the diagonal elements) representing similarity within the same subject across different sessions, and other elements representing similarity between different subjects. Bottom: within-subject between-session similarity of the first four WOCs is significantly higher than between-subject similarity (*P* < 0.001, permutation test).

We quantitatively analyzed the robustness of these components. First, we calculated similarity between the WOCs between two datasets (therefore across different electrode layouts) by spatial interpolation, and found that the first six WOCs exhibit very high similarity (absolute values of conjugate contrasts are all greater than 0.75) (**Fig. 3e**). Additionally, we performed individualized WOCCA on Dataset 2 for each experiment session of each subject and calculated average similarity of the first four WOCs between sessions and subjects. The results showed that similarity of the same subject in different sessions was significantly higher than that between two different subjects (0.877 vs. 0.808, permutation test, *P* < 0.001, also see **Supplementary Figure 4**) (**Fig. 3f**), suggesting that the four dominant alpha traveling wave patterns identified using WOCCA in resting state EEG are highly robust, and possibly contain individual information that are subject-specific.

### Differences in traveling waves between EC and EO states

To further quantify and compare the intensity and directionality of wave patterns represented by these WOCs in different states, we chose traveling energy and directionality asymmetry as statistics for describing the wave patterns. Assume we map a set of phasemaps to a target component’s projection axis using conjugate contrast function (**Fig. 4a left**), the traveling energy of this component is defined as the average squared projection values of the dataset. Greater traveling energy indicates higher intensity of this component, regardless the direction is positive or negative relative to the component (**Fig. 4a middle**). Directionality asymmetry is defined as the average of all projection values, thus positive/negative signs indicate the balance between positive and negative directions (**Fig. 4a right**).

**Figure 4:**
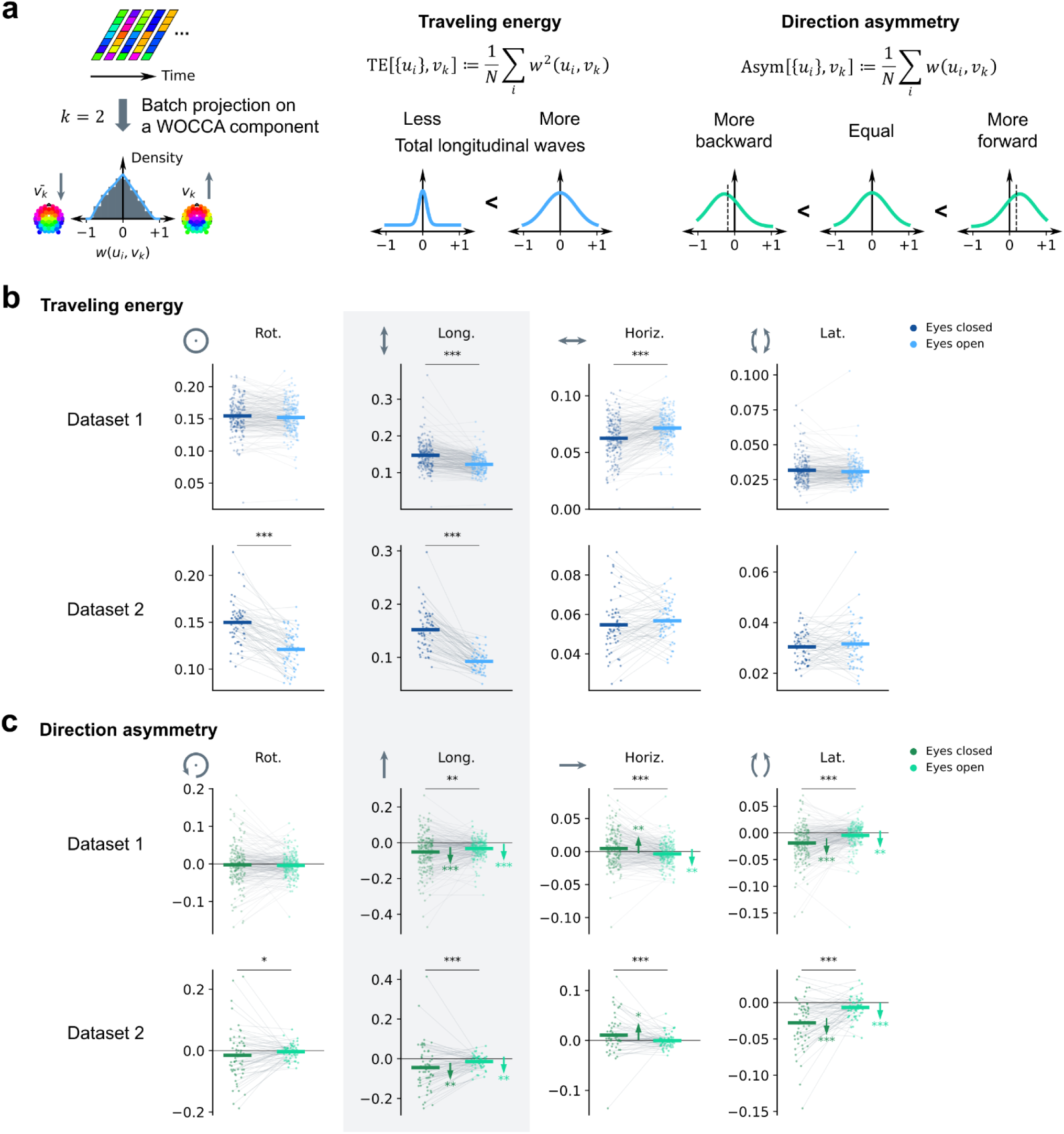
Traveling Wave Patterns and Directionality Differences between Eyes-Closed and Eyes-Open Resting States. a. Calculating the traveling wave statistics based on WOCCA. For a given WOC, the selected project phasemaps in the dataset of interest onto the component (left). The “traveling energy” represents the total presence of traveling waves belonging to the selected component in the dataset (middle), while the “directionality asymmetry” indicates dominance of a particular propagation direction of the selected component (right). b. Traveling energy statistics of the first four WOCs in EC and EO resting states for Datasets 1 and 2. Comparing EC and EO, the statistically significant result consistent in both datasets is: Component 2 (longitudinal wave) EC>EO. c. Analogous to **b**, but analyzing directionality asymmetry statistics. Comparing EC and EO, the statistically significant results consistent in both datasets are: Component 2 (longitudinal wave in forward direction) EC<EO; Component 3 (horizontal wave in rightward direction) EC>EO; Component 4 (lateral wave in forward direction) EC<EO. Additionally, arrows indicate significant deviation from 0 of directionality asymmetry, with both longitudinal and lateral waves significantly favoring the forward direction. ∗: *P* < 0.05; ∗∗: *P* < 0.01; ∗∗∗: *P* < 0.001.

We computed the traveling energy (**Fig. 4b**) and directionality asymmetry (**Fig. 4c**) statistics for the first four WOCs of Dataset 1 (for all subjects) and Dataset 2, and compared between EC and EO resting states. We noticed consistently significant results for the longitudinal wave. For both datasets, traveling energy of longitudinal wave is higher at EC than EO state (paired t test for Dataset 1, *P* = 6.0 × 10^−24^, Cohen’s *d* = 0.809; two-way ANOVA for Dataset 2, *P* = 2.1 × 10^−69^, 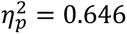), and directionality asymmetry of longitudinal wave favors more backward direction at EC than EO (for Dataset 1 *P* = 0.0024, *d* = −0.215; for Dataset 2 *P* = 6.1 × 10^−6^, 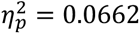). This indicates that at EC resting state during which visual input was blocked, the overall strength of alpha traveling waves in forward-backward direction was enhanced, and the balance of directionality was biased towards backward or “anterior to posterior” direction.

Directionality asymmetry of lateral wave, the other pattern that travels in forward-backward direction, also biased towards backward direction at EC state, supporting the result above. In addition, horizontal wave biased towards rightward direction at EC state for both datasets. For the other traveling wave patterns, although we didn’t observe consistently significant results for both datasets, the traveling energy of rotational wave is higher at EC state, and horizontal wave is lower at EC state. A complete summary of details is provided in

### Supplementary Table 1 and 2

Overall, the results indicate that between different brain states, there are significant differences in the dominant traveling wave patterns. In the comparison of EC and EO resting state, longitudinal and lateral wave patterns (i.e., the two forward-backward patterns) seem to play an important role: in the absence of visual information input during EC resting state, alpha traveling waves have a stronger tendency to propagate backward or “anterior to posterior” direction, compared to EO state during which visual input is present. This supports the theory of forward and backward traveling waves reflecting information transfer of bottom-up and top-down signals.

In addition, we analyzed whether directionality asymmetry of each component significantly deviates from 0 (i.e., the proportion of positive and negative wave patterns is equal). It was found that in both datasets, the direction asymmetry of longitudinal wave favors backward direction at both EC and EO state, the horizontal wave favors rightward direction at EC state, and the lateral wave also favors backward direction at EC and EO. The other components or states did not exhibit a significant effect, or the effects differ between datasets. More details are provided in **Supplementary Table 3**. Asymmetry of traveling wave directionality is common, suggesting existence of directed and unbalanced inter-areal communication supported by traveling waves.

### Traveling wave markers of diminishing consciousness

While the distinction between EC and EO states has established a preliminary connection between forward-backward traveling waves and inter-areal information flow, it is necessary to extend our findings to brain states beyond normal resting conditions. Reduced consciousness level induced by sedatives, being an important type of altered brain states, has been found to reflect disruption of bottom-up sensory information flow (**Tauber et al., 2024**). Therefore, we are interested in whether these changes during diminishing consciousness could be explained by altered alpha traveling wave patterns.

Dataset 3 (**Chennu et al., 2016**) recorded EEG data from 20 healthy participants who received certain amounts of propofol infusion during a sedative induction procedure. The procedure includes a baseline stage, a mild sedation stage with a small amount of propofol infusion, the moderate sedation stage with a medium amount of propofol infusion, and a finalizing recovery stage. The dataset also included measurements of subject behavior and propofol concentration in blood (**Figure 5a**). The behavioral task was a simple auditory discrimination task, which showed that as blood propofol level increased during the mild and moderate sedation stages, some subjects lost the ability to correctly identify auditory stimuli.

**Figure 5:**
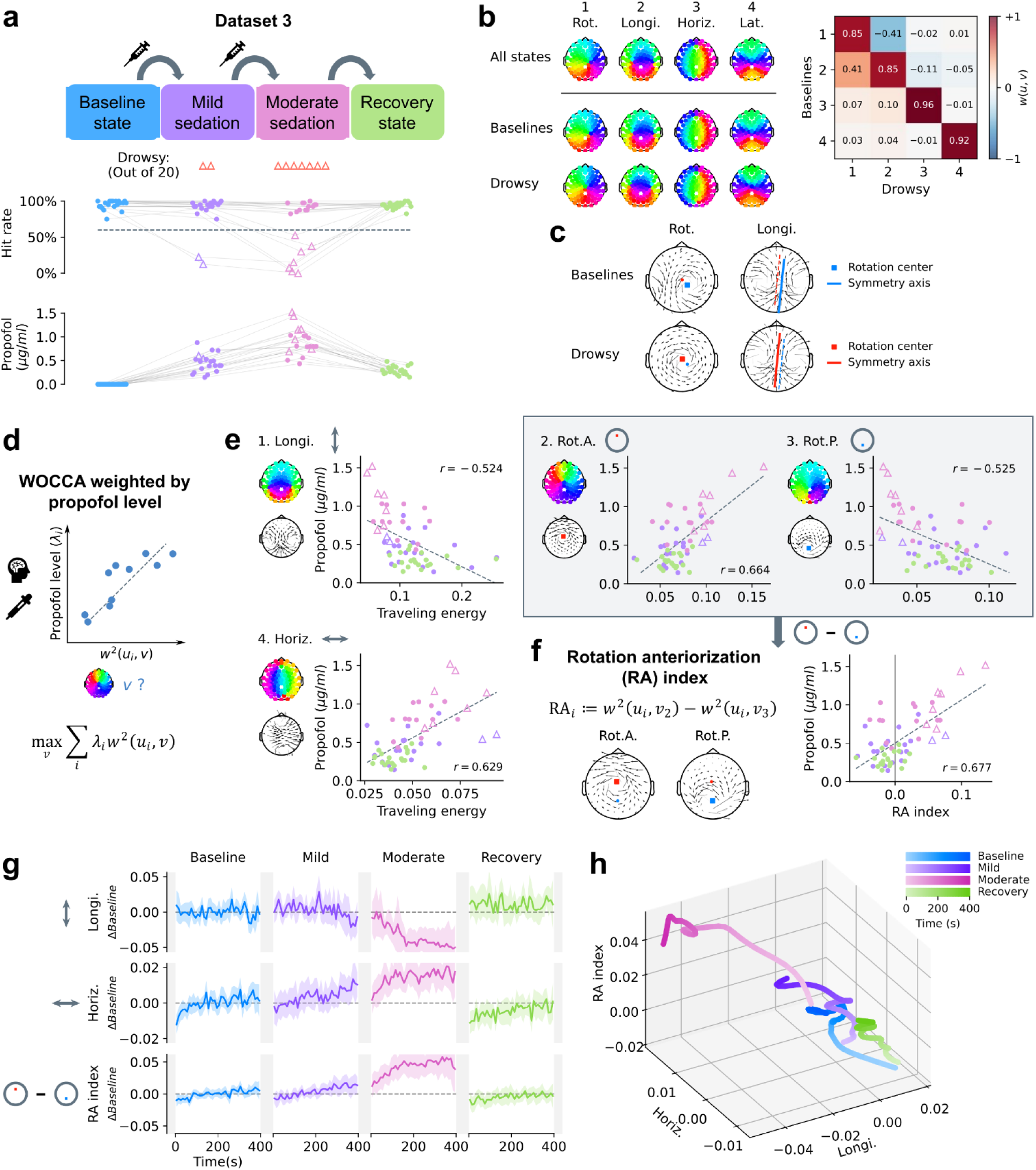
Alpha Traveling Wave Dynamics of Diminishing Consciousness. a. Experimental design, behavioral discrimination accuracy and propofol concentration in blood of subjects at all stages (more details see **Chennu et al., 2016**). Set threshold of behavioral discrimination accuracy at ≤60% to define whether a subject was in reduced consciousness (“drowsy”) state. b. The first four WOCs obtained by WOCCA from the entire dataset, baseline stage data, and the part tagged as drowsy state, respectively (left). Differences can be observed in the first (rotational) and second (longitudinal) WOCs between baseline stage and drowsy state (right). c. From the perspective of phase gradient field, the positions of rotation centers and longitudinal main axes of drowsy state deviate to the front left at drowsy state compared to baseline. d. Weighted WOCCA on propofol concentration in blood, to maximize the explanatory power of its derived traveling wave patterns for blood propofol levels. e. The first four Weighted Weakly Orthogonal Components (W-WOCs) obtained by propofol level-weighted WOCCA, along with correlations of their traveling energy statistics with the blood propofol levels. f. From the perspective of phase gradient field, the second and third wave patterns in **e** are identified as “anterior-rotational” and “posterior-rotational” waves with rotation centers shifted to anterior and posterior locations, respectively. The difference in their traveling energy is defined as the “Rotation Anteriorization Index” (RA Index). g. Traveling energy of the first (longitudinal) and fourth (horizontal) traveling wave patterns obtained by propofol level-weighted WOCCA, together with the RA index, plotted over time for each stage. Shaded areas represent *α* = 0.05 confidence intervals. h. Trajectories of traveling wave state over time for the four stages, with the three statistics in **g** serving as coordinate axes. Trajectories are smoothed using a second-order Butterworth low-pass filter with cutoff frequency of 0.01*Hz*.

Using WOCCA, we obtained sets of first four WOCs separately from the entire dataset (all stages), baseline stage (i.e., before propofol infusion), and the low consciousness level "drowsy" state data in which the test subject lost the ability to identify auditory stimuli. Dominant components obtained from the three subsets of data all exhibit patterns that are recognizable as rotational, longitudinal, horizontal, and lateral waves. However, the first two components are noticeably different between baselines and drowsy state and have lower similarity (**Fig. 5b right**, more comparisons see **Supplementary Figure 5**). We detected the positions of center singularities of rotational components and the main axes of the longitudinal components from their phase gradient fields, and found that compared to the baselines, the position of rotation center and longitudinal main axis of drowsy state deviate to the front left (**Fig. 5c**). This suggests that the dominant traveling wave patterns themselves may differ in morphological structures under different levels of consciousness.

To find the wave patterns that best reflect the differences in consciousness levels, we used blood propofol level as a quantitative measure of consciousness level and performed supervised WOCCA weighted on propofol levels to obtain the Weighted Weakly Orthogonal Components (W-WOCs) that maximally explain the differences in propofol levels between different stages (**Fig. 5d**). The first four W-WOCs are longitudinal wave, a “anterior-rotational” wave with anterior rotation center, a “posterior-rotational” wave with posterior rotation center, and horizontal wave, respectively, all with significant correlations with propofol levels (permutation test, *P* < 0.001 for all four components, also see **Supplementary Figure 6**) (**Fig. 5e**). At higher blood propofol level, traveling energy of the longitudinal wave is lower, anterior-rotational wave is higher, posterior-rotational wave is lower, and horizontal wave is higher.

In particular, from the perspective of phase gradient field, the main morphological difference between anterior- and posterior-rotational waves is indeed the positions of rotational centers (**Fig. 5f left**). We further defined the difference in traveling energy between anterior- and posterior-rotational waves as "Rotation Anteriorization" (RA) index and found this index exhibit strong positive correlation with propofol concentration, stronger than all traveling energy statistics of single wave pattern. Together, these results connected reduced consciousness states to less longitudinal waves, more horizontal waves, and more anteriorized rotational waves.

EEG in Dataset 3 was recorded continuously for approximately 7 minutes starting from the start of resting state (for baseline and recovery stages) or equilibrium of blood propofol level (for mild and moderate sedation stages), for every stage of each subject. We monitored the continuous dynamic changes of traveling wave patterns using traveling energy of longitudinal and horizontal waves, as well as the RA index, and compared them to the average level of the baseline stage (**Fig. 5g**, and a smoothed three-dimensional visualization in **Fig. 5h**). During the baseline and recovery stages, all three indices remained close to the average baseline level; during the mild sedation stage, the indices slowly trended towards the state of reduced consciousness; during the moderate sedation stage, all three indices rapidly moved in the direction of diminishing consciousness within approximately 2 minutes after start of EEG recording and then stayed in proximity to this reduced consciousness state, away from the average baseline level. This suggests that the wave patterns found by weighted WOCCA may serve as neural markers for consciousness levels, and revealed a dynamic process of rapid diminishing and then maintaining low level of consciousness during moderate sedation.

### Traveling waves explaining microstate transition statistics

Compared to the dynamic approach of WOCCA that treats EEG as segments of traveling wave patterns, classic microstate analysis assigns instantaneous snapshots of electric potential topographies to a discrete set of microstate labels. Theoretically, the characteristics of microstate sequences should be captured in the dynamic “clips” of traveling wave patterns. To investigate this relationship, we used EEG microstate analysis (**Pascual-Marqui et al., 1995**; **Michel and Koenig, 2017**) to describe alpha-band topographies in Dataset 1, and obtained four typical microstate templates (**Fig. 6a**). Meanwhile, we unfolded the first two WOCs of Dataset 1 from the complex domain into a continuous sequence of topographies along time, and assigned each of them to a microstate label (**Fig. 6b**). We noticed that the rotational and longitudinal traveling waves contain instantaneous topography patterns highly similar to microstate templates, and the order of transitions are determined by the directionality of traveling waves. For example, the positive (counterclockwise) rotational wave naturally generates a transition from microstate B to microstate A due to the rotational movement of the positions of positive and negative electric potential peaks; whereas the negative (clockwise) rotational wave generates microstate transition in the opposite order. This suggests that the traveling wave directionality may determine the transition statistics of microstate sequences, thereby reflecting the dynamic characteristics of EEG data.

**Figure 6:**
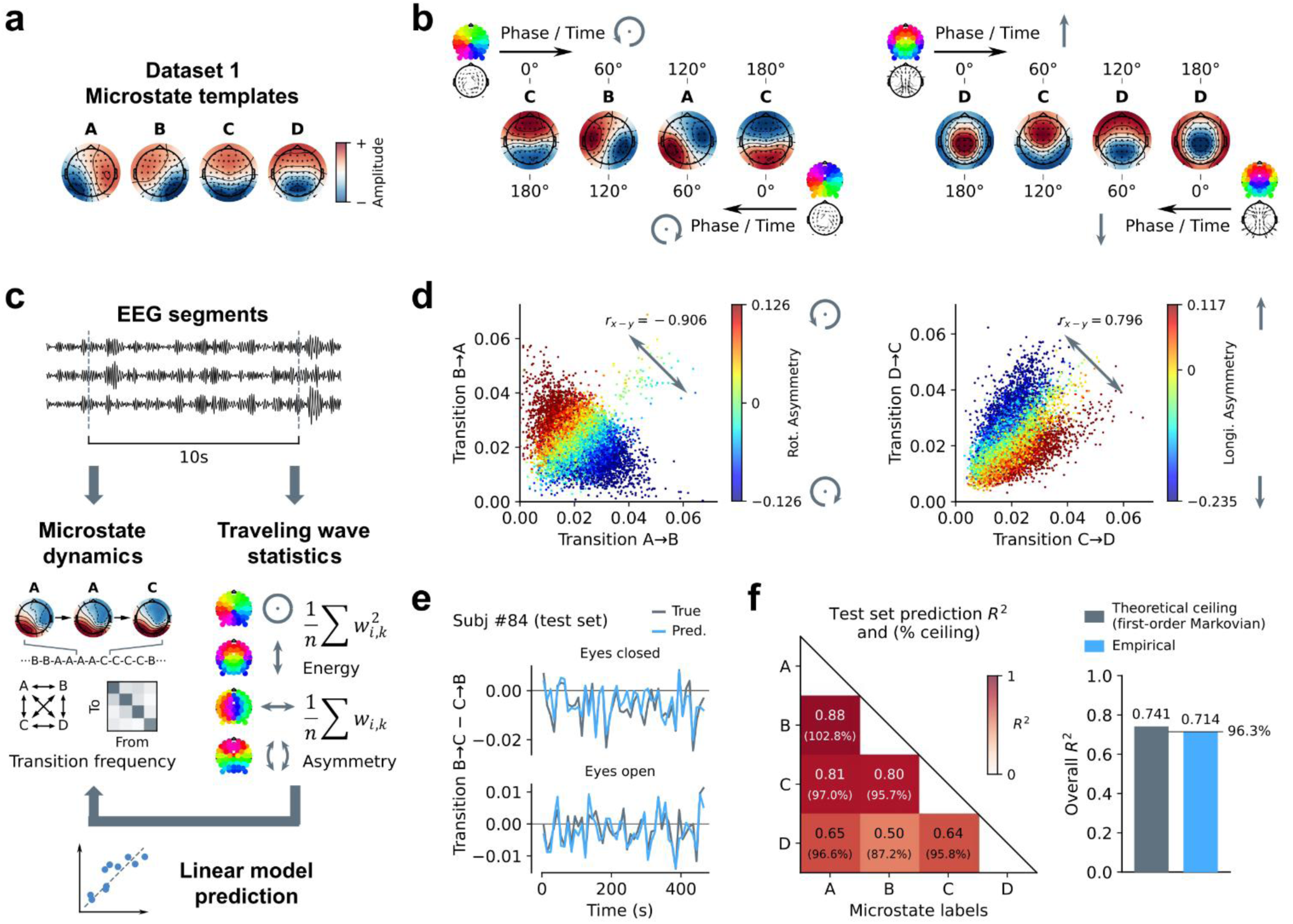
Traveling Waves Encompass Transitions in Microstate Sequences. a. Microstate templates obtained by 4-cluster analysis of resting state alpha-band EEG in Dataset 1. b. Transition dynamics of microstate sequences are governed by traveling waves: temporal unfolding of a phasemap corresponds to gradual transitions of the microstate topography, while the complex conjugate (inversion of propagation direction) of the phasemap represents reversal of the microstate sequence. c. Schematic of the analysis method: EEG signals were split into 10-second segments (top), from which the transition frequencies between four microstates were extracted as dynamical features of microstates (middle left), and traveling energy and directionality asymmetry statistics of the first four WOCs serve as static statistical features of traveling waves (middle right). Traveling wave features are then used to linearly predict microstate dynamical features. d. Left: directionality asymmetry of rotational traveling waves accurately predicts transition asymmetry between microstate A and B (i.e., the difference in frequencies of transitions from A to B and from B to A). Right: directionality asymmetry of longitudinal traveling waves accurately predicts the transition asymmetry between microstate C and D. e. Example of linear prediction of the transfer difference between microstate B and C of Subject #84, where the predicted values closely match the dynamic fluctuations of the empirical values over a recording segment. f. Left: prediction performance, measured by *R*^2^, for transition asymmetry between each pair of microstates, along with the percentage of the theoretical upper bound under first-order Markov process assumption for measuring transition probabilities from 10-second segments. Right: overall *R*^2^and its theoretical upper bound.

In light of this, we proposed a method to quantify the relationship between traveling wave patterns and transitions in microstate sequences. For a set of 10 second EEG segments, we first converted them to microstate label sequences and calculated the frequency of microstate transitions within each segment (**Lehmann et al., 2004**; **Brodbeck et al., 2012**), which is an estimate of microstate transition matrix (**Gärtner et al., 2015**) and serves as the statistics representing microstate transitions. On the other hand, we computed the traveling energy and directionality asymmetry of the first four WOCs for each segment and tried to predict the microstate transition statistics using these traveling wave features.

Consistent with our previous qualitative discussion, in real EEG data the order of transition between two microstates is closely related to the directionality of traveling waves. For example, the transition asymmetry between microstate A and B (i.e., the frequency difference, representing whether the dominant order is A to B or B to A) exhibits strong negative correlation with the directionality asymmetry of rotational waves (**Fig. 6d, left**). Similarly, for longitudinal waves and microstate C and D, the more dominant forward waves are, the more dominant is the transition from C to D (**Fig. 6d, right**).

We used the traveling wave statistics described above to predict transition asymmetry between pairs of microstates, and trained an ordinary linear regression model using the training split of Dataset 1, then evaluated model performance on the test split (**Fig. 6f**). The model predicted transition asymmetry between all six pairs of microstates with high performance, with overall *R*^2^ = 0.714. Since the amount of transition events in a 10s segment is limited, the prediction performance of the regression model is severely limited by randomness. Therefore, we calculated the theoretical variance of perfectly observed transition frequency under the assumption that microstate transitions obey a first-order Markov process, and estimated the theoretical upper bound of prediction performance accordingly. The overall empirical prediction performance of our model is *R*^2^ = 0.714, compared with the first-order Markovian upper bound of *R*^2^ = 0.741, reaching 96.3% of the ceiling level. Specifically, during the approximately 8 minutes of EC or EO resting state EEG of each subject, the fluctuation of microstate transitions across a series of 10s segments can be accurately described by our model using traveling wave statistics (example shown in **Fig. 6e**). These results indicate that natural features of traveling waves under our WOCCA framework, especially directionality asymmetry, reflects the transition statistics of microstates, and the dominant order of microstate transitions in EEG segments can be explained as rapid temporal dynamics captured by dominant traveling wave patterns.

## Discussion

Comprehending the complexity of alpha wave spatiotemporal structure and its physiological significance has long been a challenging endeavor. This study systematically examined the global spatiotemporal structure of alpha-band EEG from the perspective of traveling waves, utilizing our WOCCA method, conceived based on wave directionality, to identify the rich traveling wave patterns in alpha oscillations, including rotational, longitudinal, and lateral patterns (**Fig. 3**). The hallmark of the WOCCA method lies in its direct operation on the phasemap representation of periodic traveling waves, as well as its optimization objective of maximizing the explanatory power for wave directionality. Combined, these two features enabled WOCCA to avoid the drawbacks of CPCA and VSVD methods (**Fig. 2**), thus significantly enhanced the precision and intuitiveness of observing and quantifying traveling waves.

Identifying whole brain traveling wave propagation characteristics enables us to directly infer how alpha waves support functions such as dynamic modulation of neural activity and communication (**Muller et al., 2018**). For instance, the extensively studied forward-backward longitudinal alpha traveling waves have been closely associated with the bottom-up information from the occipital lobe through the visual pathway and top-down prior information (**Alamia and VanRullen, 2019**) or attention control signals (**Patten et al., 2012**; **Alamia et al., 2023**) from higher-level areas such as the frontal cortex. Our analysis of resting state (**Fig. 4**) reveals that overall longitudinal traveling energy is lower at EO state; while in terms of directionality, both EC and EO states are dominated by backward propagation of both longitudinal and lateral patterns, though the backward dominance is stronger at EC state, which has no visual input. We propose forward and backward alpha traveling waves are both present, with their directionality reflecting the balance of bidirectional information flow between primary and higher-level areas in the visual pathway. The presence of visual input, at a macroscopic level, competes with top-down information, manifesting as relatively weaker backward waves. This also provides a possible explanation for the observed separation of bottom-up and top-down traveling waves into high frequency (gamma) and mid-to-low frequency bands (alpha or theta) in some studies (**Van Kerkoerle et al., 2014**; **Aggarwal et al., 2022**): the dominance of backward waves and the overall reduction of forward-backward waves in presence of visual input may make it difficult to observe mid-to-low-frequency bottom-up signals, particularly when assessed using trial-averaged phase differences or gradients.

In this context, while forward-backward traveling waves potentially support communication between brain areas of multiple hierarchies, we speculate that horizontal traveling waves may facilitate two-hemispheric interactions. The high efficiency of visual information processing at EO state may rely on horizontal integration of information within the visual pathway, manifested as stronger horizontal waves. Literature on alpha oscillations and its phase coupling supporting inter-hemispheric communication agrees with our findings (**Doron et al., 2012**; **Stefanou et al., 2018**). In contrast, rotational traveling waves are more intricate, with the presence of rotational singularity indicating the absence of a consistent local propagation direction, suggesting local communication circuits being idle or suppressed. Therefore, the anteriorization of rotational singularities in a sedated state (**Fig. 5**) might signify that higher-level brain areas related to consciousness level, such as the prefrontal cortex, are probably in an inhibited state. This mechanism could be synergistic with the anteriorization of alpha oscillation power observed during mild anesthesia (**Supp et al., 2011**; **Vijayan et al., 2013**).

A widely accepted theory that explains the physiological functions of alpha waves is the "gating by inhibition" theory (**Jensen and Mazaheri, 2010**). The electric potential fluctuations of alpha oscillations lead to local neuronal population’s periodic excitation and inhibition (**Haegens et al., PNAS, 2011**), enabling the cycle period and amplitude of alpha waves to determine the rhythm and effectiveness of local information processing, respectively. This provides a powerful explanation for individuals’ temporal resolution of cognitive processing (**Surwillo, 1961**; **Samaha and Postle, 2015**) and the relationship between alpha power spatial distribution and attention selectivity (**Haegens et al., J Neurosci., 2011**; **Peylo et al., 2021**). Moreover, traveling waves provide a spatiotemporal framework for coordination of neural activity, supporting complex neural computation pathways (**Daviz et al., 2020**) or temporal order required for inter-areal communication (**Muller et al., 2016**; **Dickey et al., 2021**) by dynamic modulation of excitation and inhibition across areas. Combined with our finding that traveling wave directionality supports inter-areal communication, the above framework offers a direct interpretation of the physiological significance of traveling waves: alpha traveling waves are a mechanism for regulating large-scale brain network interactions and coordination, with the wave directionality features obtained by WOCCA representing the dominant temporal order or communication directions among brain areas. However, due to the limited spatiotemporal resolution of EEG, the precise relationship between traveling waves and the more intricate and subtle computational functions remains to be elucidated. Our results also cannot yet pinpoint the internal source or the upstream contributing brain areas of this regulatory mechanism, with some studies pointing to the importance of cortico-thalamic feedback loops (**Saalmann et al., 2012**; **Vijayan and Kopell, 2012**; **Halgren et al., 2019**), though the exact mechanisms generating and controlling alpha traveling waves remain unclear. We expect that further research in these directions will provide a more comprehensive reveal of the physiological origin of alpha traveling waves.

Traveling wave directionality also naturally encompasses the EEG microstate transitions. We found that transitions in microstate sequences are a manifestation of the propagation of traveling waves (**Fig. 6**), providing a more parsimonious representation of this phenomenon. Microstate transition frequencies have been associated with various physiological properties, such as consciousness, cognitive workload, and age (**Brodbeck et al., 2012**; **Bréchet et al., 2019**; **Zanesco et al., 2020**). Via wave directionality, the underlying causes of these physiological associations can be more directly connected to the computational and communicative functions of brain networks, while also offering insights into a wide range of microstate-related studies, particularly those investigating microstates’ source localization (**Milz et al., 2017**; **Custo et al., 2017**) and correlations with functional networks (**Britz et al., 2010**; **Panda et al., 2016**).

Lastly, while WOCCA is a powerful tool for analyzing and interpreting global traveling wave patterns, many other traveling wave analysis methods remain significant. For example, as we observed in our analysis of phase gradient fields, the orientation and distribution of rotational singularities largely determines the global wave structure (**Ermentrout and Kleinfeld, 2001**; **Townsend and Gong, 2018**; **Von Wegner, et al., 2021**), and the distribution and movement of rotational singularities in fMRI ultra-low-frequency waves have been shown to bear cognitive significance (**Xu et al., 2023**). On the other hand, simpler measures like phase differences or local phase gradients (**Ito et al., 2004**; **Liang et al., 2023**), being computationally efficient, also offer more easily interpretable metrics for local features of traveling waves. Other neuroimaging modalities like ECoG and fMRI typically possess higher spatial resolution than EEG, necessitating identification and analysis of finer spatial details; thus, in such scenarios, cross-validation among multiple traveling wave features and their interpretability may be of equal importance as global wave patterns. We expect that the WOCCA method and its derived theoretical framework of the physiological functions of traveling waves may complement and synergize with other analysis methods on local wave features, expanding applicability to a broader range of scenarios.

## Methods

Algorithms implemented for this study is programmed in Python, dependent tools and packages including MNE-Python (**Gramfort et al., 2013**), NumPy (**Harris et al., 2020**), SciPy (**Virtanen et al., 2020**), Matplotlib (**Hunter, 2007**), scikit-learn (**Pedregosa et al., 2011**), statsmodels (**Seabold and Perktold, 2010**) and PyTorch (**Paszke et al., 2019**).

### Phasemap and phase gradient vector field

For multichannel neural oscillation data including alpha-band EEG, phasemaps serve as a natural way to represent traveling wave structures. To reduce redundancy and alleviate the impact of noise as well as undefined phases at wave nodes, we adopted an indirect approach of principal eigenvector method, instead of directly employing the Hilbert transform, to extract phasemaps from neural oscillation data. First, the data is divided into reasonably sized segments bounded by Global Field Power (GFP) peaks (**Lehmann and Skrandies, 1980**) that appear at approximately double the frequency of the ongoing oscillation, with *u*_*i*_ denoting the electric potential at channel *i* at a given time point:

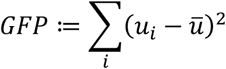

Subsequently, for Hilbert transformed analytical signals ℎ_*i*_, we calculated complex coherence between every pair of channels *i*, *j*, yielding a complex coherence matrix, whose phases and magnitudes capture the average phase difference and strength of synchronization between each pair of channels, respectively:

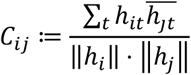

Matrix *C* can be diagonalized since it’s Hermitian. By computing the principal eigenvector (following the approach in **Deco et al., 2019**, but applied to complex-valued matrices), its phases represent the dominant phase pattern of the segment, while its magnitudes represent the temporal consistency of phase differences between each channel and all other channels, providing a natural "reliability" weighting for each channel’s phase. This method also naturally normalizes the magnitude of each phasemap to 1.

To compute phase gradient vector fields from phasemaps, we first defined the spatial neighbors for each channel. In a synthetic grid setting, up to 8 directly and diagonally adjacent channels are naturally chosen as the neighbors. For irregular EEG electrode layout, we applied Delaunay triangulation (implemented in MNE-Python) to define neighbors. The phase differences between a channel and its neighbors were modeled as a linear function of vectors connecting their spatial positions, and the sum of squared errors were minimized through gradient descent (the classic linear least square method cannot be used due to periodicity of phase), where *p*_*i*_ ≔ [*x*_*i*_, *y*_*i*_] denotes the spatial position of channel*i*, *N*(*k*) denotes the set of neighbors of channel *k*, and 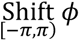 denotes shifting *ϕ* to an equivalent phase *ϕ* + 2*kπ* (*k* ∈ ℤ) within interval [−*π*, *π*):

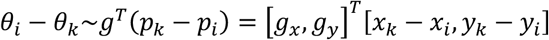

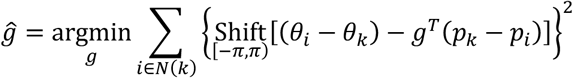

In this way, the estimated phase gradient vector *g*^ represents the direction from leading to lagging phase within the neighbors of the selected channel. Additionally, the fit’s *R*^2^ can be calculated using the final sum of squared errors. Higher *R*^2^indicates a more stable spatial linear distribution of phase, thus a more regular local traveling wave; and vice versa.

### Implementation of the WOCCA algorithm

Implementing WOCCA involves constrained nonlinear optimization algorithms. If naïvely following WOCCA’s original formulation, computing the optimization objective would require going through the whole phasemap dataset for every iteration. For a more efficient implementation, we transformed the phasemaps into skew-symmetric matrices:

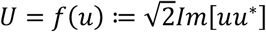

In **Supplementary Note 1**, we proved that the linear superposition of the skew-symmetric matrices *U* is equivalent to the nonlinear combination of phasemaps *u* in terms of traveling wave patterns, with the conjugate contrast function being equivalent to the element-wise inner product of their transformed skew-symmetric matrices. Therefore, we can directly optimize over the set of transformed skew-symmetric matrices:

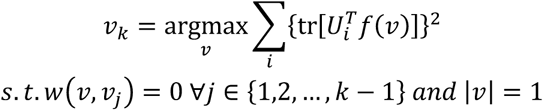

Since the objective function is now a quadratic form of *f*(*v*), precomputing the quadratic form matrix avoids repeatedly going through the whole phasemap dataset during iterations, which greatly improves computational efficiency. Finally, we employ a constrained nonlinear optimization algorithm (specifically, the trust-constr algorithm implemented in SciPy) to solve the optimization problem.

Furthermore, due to the low efficiency and convergence difficulties of high-dimensional nonlinear optimizations, for real EEG data, we performed preliminary dimensionality reduction (pre-reduction) on the phasemap data before WOCCA. Specifically, we conducted real-valued Principal Component Analysis (PCA) on both the real and imaginary parts of the phasemaps, and retained the top 30 spatial principal components. The phasemaps are then projected onto this 30-dimensional subspace before applying WOCCA. The resulting WOCs in the subspace are subsequently reprojected back into the original EEG channel space using the principal components, yielding the final decomposition result. This approach’s validity rests on: first, orthogonal transformations do not affect inner products and thus do not alter the values of conjugate contrast functions; second, since the conjugate contrast function is the difference of squared magnitudes of two inner products, if dominant traveling wave patterns exist, the WOCs with larger traveling energy will be captured by the leading real-valued principal components (though the converse may not hold). See **Supplementary Figure 3** for comparison of the results and efficiency between direct and pre-reduced WOCCA on a synthetic dataset.

### The synthetic phasemap dataset

We establish a setup of uniform 6×6 grid on a virtual unit-square canvas. Both rotational and longitudinal phasemaps were constructed on the grid, designing only the phases while keeping magnitude of all channels at 1/6 (ensuring the magnitude of phasemaps equals 1). For the rotational waves, given a rotation center position *O* ≔ [*x*_0_, *y*_0_] and a winding number *n*_*W*_ ∈ ℤ, the phase at each channel location *P*_*i*_ ≔ [*x*_*i*_, *y*_*i*_] is set as a multiple of the angle between the line connecting *P*_*i*_ to *O* and the positive x-axis:

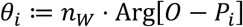

For the longitudinal waves, given a propagation direction angle *ϕ* and a wavelength *λ*, the phase at position *P* is set as a multiple of its projection along the propagation direction:

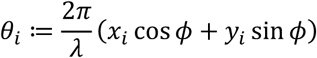

Our synthetic dataset consists of 6,000 randomly constructed phase patterns, including 2,000 rotational wave patterns, whose rotation centers obey a bivariate normal distribution with mean of [0.5, 0.5], independent standard deviations of 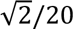 along both axes, split evenly between 1,000 instances with a winding number of +1 (counterclockwise rotation) and 1,000 with −1 (clockwise rotation). The remaining 4,000 longitudinal wave patterns have propagation direction angle *ψ* + *δ*, in which *ψ* was set at +*π*/2 (forward propagation) and −*π*/2 (backward propagation) for 2,000 instances each, and *δ* was drawn from a normal distribution with mean of 0 and standard deviation of*π*/12. The wavelength *λ* is uniformly distributed between 0.5 and 3.

### EEG datasets and preprocessing

This study employs three publicly available EEG datasets from independent sources, with Datasets 1 and 2 providing resting-state EEG data, and Dataset 3 providing EEG data at sedated state induced by propofol infusion. Below is a brief overview of the datasets; for more comprehensive details, refer to the original publications of each dataset.

Dataset 1, the Mind-Brain-Body dataset (**Babayan et al., 2019**), contains resting-state EEG, MRI, and additional physiological and psychological metrics from a total of 227 healthy participants across various age groups. The resting-state EEG protocol consists of alternating 1-minute epochs of eyes-closed and eyes-open resting states, repeated eight times for a total recording duration of 16 minutes. EEG signals were captured using the extended 10-20 system across 61 valid channels. We selected 203 subjects with complete post-preprocessed EEG data (138 in young group aged 20-40, 65 in elderly group aged 55-80; 74 females, 129 males). Among these, data from 83 subjects without any channels marked as “bad” (i.e., all EEG channels were usable) were assigned to the training split (61 young; 23 females), and data from the remaining 120 subjects were assigned to the test split (77 young; 51 females). Dataset 2, the Test-Retest Resting and Cognitive State EEG dataset (**Wang et al., 2022**), contains EEG at resting state and self-paced light cognition tasks, as well as behavioral and other physiological/psychological metrics, from 60 healthy young participants (aged 18-28, mean age 20.01 years; 32 females, 28 males) across three experimental sessions. Each session involved continuous 5-minute blocks of eyes-closed rest, eyes-open rest, self-paced arithmetic, self-paced memory recall, and singing a song within one’s head. Following the first session of each subject, a second session was carried out approximately 90 minutes later, and a third one about one month later, with identical procedures. EEG signals were captured using the extended 10-20 system across 61 valid channels, and no bad channels were reported in the original publication.

Dataset 3, Brain Connectivity during Propofol Sedation (**Chennu et al., 2016**), contains continuous EEG, behavioral data, and blood propofol concentration measurements from 22 healthy volunteers before and after propofol infusion. The experiment consisted of four stages: baseline (prior to propofol infusion), mild sedation (target plasma concentration 0.6*μg*/*ml*), moderate sedation ( 1.2*μg*/*ml*, ideally the subjects get fully relaxed but still possibly responsive to tasks), and recovery (20 minutes after moderate sedation stage). Approximately 7 minutes of eyes-closed resting state EEG was recorded during each stage, followed by a simple auditory discrimination task to assess accuracy and reaction time. Blood samples for propofol concentration were taken in the latter three stages. EEG signals were captured using a high-density 128-channel system, with 91 valid channels on scalp. The original publication selected 20 subjects with intact EEG data (mean age 30.85 years; 11 females, 9 males) and interpolated over bad channels identified during preprocessing.

Further preprocessing of the EEG data was applied to each dataset, including resampling to 250*Hz*, filtering to the alpha frequency range of 7-13*Hz*, and applying average referencing. Subsequent extraction of phasemaps and phase gradient fields were based on this further preprocessed data.

### Statistical methods for the stability of WOCs

When comparing the similarity of WOCs between Dataset 1 and Dataset 2, we employed spatial interpolation to address the discrepancy in electrode layouts between the two EEG datasets. Specifically, we transferred the weakly orthogonal components based on Dataset 1, using spline interpolation (implemented in MNE-Python) to map them onto the electrode layout of Dataset 2, and calculated absolute value of the conjugate difference function between them and the corresponding WOCs of Dataset 2. Conversely, we also transferred Dataset 2 to match Dataset 1’s layout and calculated the corresponding results. The average of these two results served as the cross-dataset similarity measure for WOCs.

When comparing similarity between different sessions of the same subject against similarity between two different subjects of Dataset 2, we performed WOCCA separately on phasemap data of each session for every subject, yielding the first 10 WOCs per session. We then matched the 10 WOCs of each session to the first four dominant WOCs of the whole Dataset 2 according to maximizing absolute values of conjugate contrast function, and obtained the four most typical WOCs of each session. For each pair of different sessions, we computed similarities of their four most typical WOCs by average of absolute conjugate contrast function, thereby constructed a similarity matrix. The matrix is symmetric by definition and we only consider its lower triangular part. To evaluate whether there was a significant difference in traveling wave pattern similarity within subjects versus between subjects, we compared within-subject similarity (the 3×3 diagonal blocks excluding the diagonal elements of the similarity matrix) and between-subject similarity (all elements outside the diagonal blocks), and conducted a non-parametric test using the Wilcoxon rank-sum statistic. We generated a null distribution by randomly shuffling subject identities and session orders 1000 times and calculating the statistics. The empirical statistic was compared to the null distribution to estimate the P-value. More details are provided in **Supplementary Figure 4**.

### Statistical methods for comparing EC and EO resting states

For a given set of phasemaps and a weakly orthogonal component, we employ traveling energy and directionality asymmetry metrics, derived from the projection onto this component, as the primary features of this dataset viewed from perspective of this traveling wave pattern. Traveling energy is defined as mean of squared conjugate contrast between the phasemap dataset and the WOC. Assuming the magnitude of phasemaps is normalized to 1, the range of traveling energy is [0,1]:

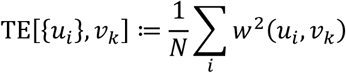

Directionality asymmetry is defined as the direct average of conjugate contrast between the phasemap dataset and the WOC. Assuming the magnitude of phasemaps is normalized to 1, its range is [−1,1]:

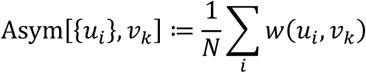

When comparing the metrics of EC versus EO resting states, to Dataset 1 we applied paired t-tests directly and calculated P-values as well as Cohen’s d effect size. For Dataset 2, given the presence of both subject and session variables, we employed two-way ANOVA that incorporates both subject identity and EC/EO state labels, in order to control the effect of subject-level individual differences. P-values and partial *η*^2^ effect sizes for the EC/EO state variable were calculated.

When examining whether there was significant directionality asymmetry, we conducted a one-sample t-test comparing directionality asymmetry statistics to 0, and calculated P-values and Cohen’s d effect size.

### Relating traveling waves to blood propofol concentrations

For Dataset 3, we applied WOCCA separately to the entire dataset (all stages of all subjects), baseline stage data (the baseline phase for all subjects), and the low consciousness level "drowsy" state data (where auditory discrimination accuracy was lower than 60%) to obtain their first four dominant WOCs. We assessed the similarity between the dominant traveling wave patterns from these three sets of WOCs by computing pairwise conjugate contrast between the phasemaps.

Regarding rotational and longitudinal traveling waves, we characterized their morphological characteristics by the position of rotation center and the orientation as well as location of the longitudinal main axis, respectively. Based on the phase gradient field, the projection length of the phase gradient vector along the direction perpendicular to the line connecting the current channel location to the rotation center should match the winding number of the rotation. Given winding number *n*_*W*_ and a phase gradient field *G* derived from a phasemap with *R*^2^ value of each channel’s phase gradient vector, we locate the rotation center *q* ≔ [*q*_*x*_, *q*_*y*_] by minimizing, via gradient descent, the sum of squared difference between the actual angular velocity calculated from projection of phase gradients and that defined by winding number. This objective function is also weighted (weights *w*_*i*_) by each channel’s *R*^2^ value and its distance to the rotation center for numerical stability:

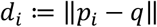

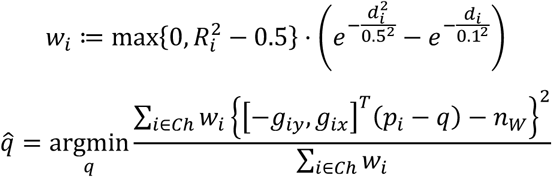

For longitudinal waves, we define the main axis as the symmetry axis between two opposing local rotational patterns present on the left and right side. For rotational waves, we set *n*_*W*_ to 1 and initialized at (0,0), approximately at the center of EEG electrode layout; for longitudinal waves, we searched for the left and right local rotation patterns with *n*_*W*_ set to 1 and −1, both initialized at (0,0).

To account for differences in blood propofol concentrations and consciousness levels, supervised weighted WOCCA maximizes the explanatory power of the weighted weak orthogonal components for current blood propofol concentration *λ*_*i*_. Specifically, the dot product of *λ*_*i*_ and the squared conjugate contrast of phasemaps with the W-WOC (i.e., traveling energy) is maximized:

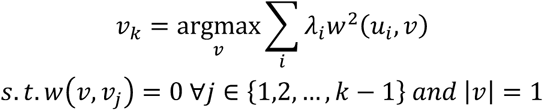

We then projected phasemaps onto these four W-WOCs for each stage of each subject and computed the traveling energy. Pearson correlation coefficients were calculated between traveling energy and blood propofol concentrations (baseline stage excluded since propofol level is always 0). We generated a null distribution by randomly shuffling blood propofol concentrations 1000 times, applied supervised weighted WOCCA, and calculated the correlation coefficients. The empirical coefficient was compared to the null distribution as a nonparametric test and P-value was estimated. More details are provided in **Supplementary Figure 6**.

When evaluating the temporal evolution of the three traveling wave indices, we computed the indices over 10-second segments and averaged across all subjects before subtracting the overall average value of baseline stage. In order to improve visibility in 3D visualization, a 0.01 *Hz* second-order Butterworth low-pass filter was applied for smoothing.

### Microstate analysis

Our approach to extracting microstate templates from continuous alpha-band EEG adheres to common practices in the literature (**Bréchet et al., 2019**; **Mishra et al., 2020**; **Wegner et al., 2021**), with minor modifications. We started from extracting electric potential topographies at GFP peaks from the alpha-band (7-13*Hz*) EEG data in Dataset 1. Next, from the training split of Dataset 1, we sampled a total of 20,000 topographies evenly from EC and EO states of each subject. We compute normalized dot products between each pair of topographies, serving as the affinity matrix, and employed spectral clustering (implemented in SciPy) with *k* = 4 to group the samples into four preliminary clusters:

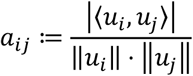

For each cluster, the first principal component vector obtained by singular value decomposition serves as the cluster centroid, yielding a candidate set of four microstate templates. Topographies are then assigned their preliminary microstate labels according to maximum similarity. This above process of randomly sampling and clustering microstates is repeated 1,000 times, and the candidate set of templates and labels that minimizes the sum of distances between each topography and its corresponding preliminary microstate template is selected as the final result of microstate analysis.

Microstate labels were assigned to the alpha-band EEG (to every sample at 250*Hz* sample rate, not just GFP peaks). Also, when analyzing microstates within a traveling wave segment represented by a phasemap *v*, we temporally unfolded the phasemap and assigned microstate labels to every phase:

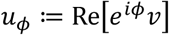

Microstate sequences were divided into 10-second segments, within which the transition frequency matrices between the four microstate labels were computed. As for traveling waves, the traveling energy and directionality asymmetry statistics of the first four WOCs of Dataset 1 are calculated for these segments.

As for assessing the relationship between microstate transition asymmetry (transition frequency from state X to Y minus Y to X) and traveling waves, we constructed a linear regression model predicting transition asymmetry using traveling wave statistics on the training split of Dataset 1, and evaluate the model’s prediction performance using *R*^2^ on the test split. To estimate the theoretical upper bound of prediction performance, for each 10s segment’s transitions from X to Y, we first calculated the number of samples in state X *n*_*X*_ and the conditional frequency *q*_*XY*_ of transitioning to Y at the next sample given the current state is X. The empirically observed number of transitions *n*_*XY*_ was modeled as a binomial distribution*B*(*n*_*X*_, *q*_*XY*_), and the variance of transition frequencies was estimated (assuming the total number of samples in a segment is *n*):

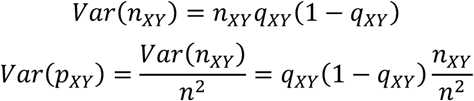

This yields the part of variance in transition asymmetry unexplained by observing this 10s segment, therefore the theoretical upper bound for *R*^2^ can be estimated accordingly:

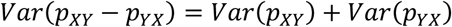

The final overall *R*^2^, both the empirical value and its theoretical upper bound, were obtained by averaging across the six pairs of microstates.

## Code and data availability

The source code of our implementation of the WOCCA algorithm, analysis of EEG datasets and creation of figures is available at https://github.com/HongLabTHU/WOCCA.

For public access of the three EEG datasets, please refer to their original publications (**Babayan et al., 2019**; **Wang et al., 2022**; **Chennu et al., 2016**).

## Acknowledgements

We thank Jingling Qu, Ruwei Yao, Xintong Yao and Diya Yi for their valuable suggestions and feedbacks.

## Supplementary information

Supplementary material is available at https://github.com/HongLabTHU/WOCCA.

## References

Berger, H. (1929). Über das elektroenkephalogramm des menschen. Archiv für psychiatrie und nervenkrankheiten, 87(1), 527–570.

Adrian, E. D., & Matthews, B. H. (1934). The Berger rhythm: potential changes from the occipital lobes in man. Brain, 57(4), 355–385.

Jokisch, D., & Jensen, O. (2007). Modulation of gamma and alpha activity during a working memory task engaging the dorsal or ventral stream. Journal of Neuroscience, 27(12), 3244–3251.

Haegens, S., Händel, B. F., & Jensen, O. (2011). Top-down controlled alpha band activity in somatosensory areas determines behavioral performance in a discrimination task. Journal of Neuroscience, 31(14), 5197–5204.

Mathewson, K. E., Gratton, G., Fabiani, M., Beck, D. M., & Ro, T. (2009). To see or not to see: prestimulus α phase predicts visual awareness. Journal of Neuroscience, 29(9), 2725–2732.

Hansen, N. E., Harel, A., Iyer, N., Simpson, B. D., & Wisniewski, M. G. (2019). Pre-stimulus brain state predicts auditory pattern identification accuracy. NeuroImage, 199, 512–520.

Schroeder, C. E., & Lakatos, P. (2009). Low-frequency neuronal oscillations as instruments of sensory selection. Trends in neurosciences, 32(1), 9–18.

Jensen, O., & Mazaheri, A. (2010). Shaping functional architecture by oscillatory alpha activity: gating by inhibition. Frontiers in human neuroscience, 4, 186.

Pascual-Marqui, R. D., Michel, C. M., & Lehmann, D. (1995). Segmentation of brain electrical activity into microstates: model estimation and validation. IEEE Transactions on Biomedical Engineering, 42(7), 658–665.

Lehmann, D., Ozaki, H., & Pál, I. (1987). EEG alpha map series: brain micro-states by space-oriented adaptive segmentation. Electroencephalography and clinical neurophysiology, 67(3), 271–288.

Michel, C. M., & Koenig, T. (2018). EEG microstates as a tool for studying the temporal dynamics of whole-brain neuronal networks: a review. Neuroimage, 180, 577–593.

Bréchet, L., Brunet, D., Birot, G., Gruetter, R., Michel, C. M., & Jorge, J. (2019). Capturing the spatiotemporal dynamics of self-generated, task-initiated thoughts with EEG and fMRI. Neuroimage, 194, 82–92.

Zanesco, A. P., King, B. G., Skwara, A. C., & Saron, C. D. (2020). Within and between-person correlates of the temporal dynamics of resting EEG microstates. NeuroImage, 211, 116631.

da Cruz, J. R., Favrod, O., Roinishvili, M., Chkonia, E., Brand, A., Mohr, C.,… & Herzog, M. H. (2020). EEG microstates are a candidate endophenotype for schizophrenia. Nature communications, 11(1), 3089.

Gui, P., Jiang, Y., Zang, D., Qi, Z., Tan, J., Tanigawa, H.,… & Wang, L. (2020). Assessing the depth of language processing in patients with disorders of consciousness. Nature neuroscience, 23(6), 761–770.

Mishra, A., Englitz, B., & Cohen, M. X. (2020). EEG microstates as a continuous phenomenon. Neuroimage, 208, 116454.

Nunez, P. L. (1974). Wavelike properties of the alpha rhythm. IEEE Transactions on Biomedical Engineering, (6), 473–482.

Townsend, R. G., & Gong, P. (2018). Detection and analysis of spatiotemporal patterns in brain activity. PLoS computational biology, 14(12), e1006643.

Aggarwal, A., Brennan, C., Luo, J., Chung, H., Contreras, D., Kelz, M. B., & Proekt, A. (2022). Visual evoked feedforward–feedback traveling waves organize neural activity across the cortical hierarchy in mice. Nature Communications, 13(1), 4754.

Liang, Y., Liang, J., Song, C., Liu, M., Knöpfel, T., Gong, P., & Zhou, C. (2023). Complexity of cortical wave patterns of the wake mouse cortex. Nature Communications, 14(1), 1434.

Davis, Z. W., Muller, L., Martinez-Trujillo, J., Sejnowski, T., & Reynolds, J. H. (2020). Spontaneous travelling cortical waves gate perception in behaving primates. Nature, 587(7834), 432–436.

Dickey, C. W., Sargsyan, A., Madsen, J. R., Eskandar, E. N., Cash, S. S., & Halgren, E. (2021). Travelling spindles create necessary conditions for spike-timing-dependent plasticity in humans. Nature communications, 12(1), 1027.

Sauseng, P., Klimesch, W., Doppelmayr, M., Pecherstorfer, T., Freunberger, R., & Hanslmayr, S. (2005). EEG alpha synchronization and functional coupling during top-down processing in a working memory task. Human brain mapping, 26(2), 148–155.

Fellinger, R., Gruber, W., Zauner, A., Freunberger, R., & Klimesch, W. (2012). Evoked traveling alpha waves predict visual-semantic categorization-speed. NeuroImage, 59(4), 3379–3388.

Patten, T. M., Rennie, C. J., Robinson, P. A., & Gong, P. (2012). Human cortical traveling waves: dynamical properties and correlations with responses. PLoS One, 7(6), e38392.

Alamia, A., & VanRullen, R. (2019). Alpha oscillations and traveling waves: Signatures of predictive coding?. PLoS Biology, 17(10), e3000487.

Alamia, A., Terral, L., D’ambra, M. R., & VanRullen, R. (2023). Distinct roles of forward and backward alpha-band waves in spatial visual attention. Elife, 12, e85035.

Klimesch, W., Sauseng, P., & Hanslmayr, S. (2007). EEG alpha oscillations: the inhibition– timing hypothesis. Brain research reviews, 53(1), 63–88.

Bolt, T., Nomi, J. S., Bzdok, D., Salas, J. A., Chang, C., Thomas Yeo, B. T.,… & Keilholz, S. D. (2022). A parsimonious description of global functional brain organization in three spatiotemporal patterns. Nature Neuroscience, 25(8), 1093–1103.

Gabay, N. C., Babaie-Janvier, T., & Robinson, P. A. (2018). Dynamics of cortical activity eigenmodes including standing, traveling, and rotating waves. Physical Review E, 98(4), 042413.

von Wegner, F., Bauer, S., Rosenow, F., Triesch, J., & Laufs, H. (2021). EEG microstate periodicity explained by rotating phase patterns of resting-state alpha oscillations. Neuroimage, 224, 117372.

Babayan, A., Erbey, M., Kumral, D., Reinelt, J. D., Reiter, A. M., Röbbig, J.,… & Villringer, A. (2019). A mind-brain-body dataset of MRI, EEG, cognition, emotion, and peripheral physiology in young and old adults. Scientific data, 6(1), 1–21.

Wang, Y., Duan, W., Dong, D., Ding, L., & Lei, X. (2022). A test-retest resting, and cognitive state EEG dataset during multiple subject-driven states. Scientific Data, 9(1), 566.

Chennu, S., O’Connor, S., Adapa, R., Menon, D. K., & Bekinschtein, T. A. (2016). Brain connectivity dissociates responsiveness from drug exposure during propofol-induced transitions of consciousness. PLoS computational biology, 12(1), e1004669.

Tauber, J. M., Brincat, S. L., Stephen, E. P., Donoghue, J. A., Kozachkov, L., Brown, E. N., & Miller, E. K. (2024). Propofol-mediated Unconsciousness Disrupts Progression of Sensory Signals through the Cortical Hierarchy. Journal of Cognitive Neuroscience, 36(2), 394–413.

Lehmann, D., Faber, P. L., Galderisi, S., Herrmann, W. M., Kinoshita, T., Koukkou, M.,… & Koenig, T. (2005). EEG microstate duration and syntax in acute, medication-naive, first-episode schizophrenia: a multi-center study. Psychiatry Research: Neuroimaging, 138(2), 141–156.

Brodbeck, V., Kuhn, A., von Wegner, F., Morzelewski, A., Tagliazucchi, E., Borisov, S.,… & Laufs, H. (2012). EEG microstates of wakefulness and NREM sleep. Neuroimage, 62(3), 2129–2139.

Gärtner, M., Brodbeck, V., Laufs, H., & Schneider, G. (2015). A stochastic model for EEG microstate sequence analysis. Neuroimage, 104, 199–208.

Muller, L., Chavane, F., Reynolds, J., & Sejnowski, T. J. (2018). Cortical travelling waves: mechanisms and computational principles. Nature Reviews Neuroscience, 19(5), 255–268.

Van Kerkoerle, T., Self, M. W., Dagnino, B., Gariel-Mathis, M. A., Poort, J., Van Der Togt, C., & Roelfsema, P. R. (2014). Alpha and gamma oscillations characterize feedback and feedforward processing in monkey visual cortex. Proceedings of the National Academy of Sciences, 111(40), 14332–14341.

Doron, K. W., Bassett, D. S., & Gazzaniga, M. S. (2012). Dynamic network structure of interhemispheric coordination. Proceedings of the National Academy of Sciences, 109(46), 18661–18668.

Stefanou, M. I., Desideri, D., Belardinelli, P., Zrenner, C., & Ziemann, U. (2018). Phase synchronicity of μ-rhythm determines efficacy of interhemispheric communication between human motor cortices. Journal of Neuroscience, 38(49), 10525–10534.

Supp, G. G., Siegel, M., Hipp, J. F., & Engel, A. K. (2011). Cortical hypersynchrony predicts breakdown of sensory processing during loss of consciousness. Current biology, 21(23), 1988–1993.

Vijayan, S., Ching, S., Purdon, P. L., Brown, E. N., & Kopell, N. J. (2013). Thalamocortical mechanisms for the anteriorization of alpha rhythms during propofol-induced unconsciousness. Journal of Neuroscience, 33(27), 11070–11075.

Haegens, S., Nácher, V., Luna, R., Romo, R., & Jensen, O. (2011). α-Oscillations in the monkey sensorimotor network influence discrimination performance by rhythmical inhibition of neuronal spiking. Proceedings of the National Academy of Sciences, 108(48), 19377–19382.

Surwillo, W. W. (1961). Frequency of the ‘alpha’rhythm, reaction time and age. Nature, 191(4790), 823–824.

Samaha, J., & Postle, B. R. (2015). The speed of alpha-band oscillations predicts the temporal resolution of visual perception. Current Biology, 25(22), 2985–2990.

Peylo, C., Hilla, Y., & Sauseng, P. (2021). Cause or consequence? Alpha oscillations in visuospatial attention. Trends in Neurosciences, 44(9), 705–713.

Muller, L., Piantoni, G., Koller, D., Cash, S. S., Halgren, E., & Sejnowski, T. J. (2016). Rotating waves during human sleep spindles organize global patterns of activity that repeat precisely through the night. Elife, 5, e17267.

Saalmann, Y. B., Pinsk, M. A., Wang, L., Li, X., & Kastner, S. (2012). The pulvinar regulates information transmission between cortical areas based on attention demands. science, 337(6095), 753–756.

Vijayan, S., & Kopell, N. J. (2012). Thalamic model of awake alpha oscillations and implications for stimulus processing. Proceedings of the National Academy of Sciences, 109(45), 18553–18558.

Halgren, M., Ulbert, I., Bastuji, H., Fabó, D., Erő ss, L., Rey, M.,… & Cash, S. S. (2019). The generation and propagation of the human alpha rhythm. Proceedings of the National Academy of Sciences, 116(47), 23772–23782.

Milz, P., Pascual-Marqui, R. D., Achermann, P., Kochi, K., & Faber, P. L. (2017). The EEG microstate topography is predominantly determined by intracortical sources in the alpha band. Neuroimage, 162, 353–361.

Custo, A., Van De Ville, D., Wells, W. M., Tomescu, M. I., Brunet, D., & Michel, C. M. (2017). Electroencephalographic resting-state networks: source localization of microstates. Brain connectivity, 7(10), 671–682.

Britz, J., Van De Ville, D., & Michel, C. M. (2010). BOLD correlates of EEG topography reveal rapid resting-state network dynamics. Neuroimage, 52(4), 1162–1170.

Panda, R., Bharath, R. D., Upadhyay, N., Mangalore, S., Chennu, S., & Rao, S. L. (2016). Temporal dynamics of the default mode network characterize meditation-induced alterations in consciousness. Frontiers in human neuroscience, 10, 372.

Ermentrout, G. B., & Kleinfeld, D. (2001). Traveling electrical waves in cortex: insights from phase dynamics and speculation on a computational role. Neuron, 29(1), 33–44.

Xu, Y., Long, X., Feng, J., & Gong, P. (2023). Interacting spiral wave patterns underlie complex brain dynamics and are related to cognitive processing. Nature human behaviour, 7(7), 1196–1215.

Ito, J., Nikolaev, A. R., & Leeuwen, C. V. (2005). Spatial and temporal structure of phase synchronization of spontaneous alpha EEG activity. Biological cybernetics, 92(1), 54–60.

Gramfort, A., Luessi, M., Larson, E., Engemann, D. A., Strohmeier, D., Brodbeck, C.,… & Hämäläinen, M. (2013). MEG and EEG data analysis with MNE-Python. Frontiers in neuroscience, 7, 70133.

Harris, C. R., Millman, K. J., Van Der Walt, S. J., Gommers, R., Virtanen, P., Cournapeau, D.,… & Oliphant, T. E. (2020). Array programming with NumPy. Nature, 585(7825), 357–362.

Virtanen, P., Gommers, R., Oliphant, T. E., Haberland, M., Reddy, T., Cournapeau, D.,… & Van Mulbregt, P. (2020). SciPy 1.0: fundamental algorithms for scientific computing in Python. Nature methods, 17(3), 261–272.

Hunter, J. D. (2007). Matplotlib: A 2D graphics environment. Computing in science & engineering, 9(03), 90–95.

Pedregosa, F., Varoquaux, G., Gramfort, A., Michel, V., Thirion, B., Grisel, O.,… & Duchesnay, É. (2011). Scikit-learn: Machine learning in Python. the Journal of machine Learning research, 12, 2825–2830.

Seabold, S., & Perktold, J. (2010). Statsmodels: econometric and statistical modeling with python. SciPy, 7, 1.

Paszke, A., Gross, S., Massa, F., Lerer, A., Bradbury, J., Chanan, G.,… & Chintala, S. (2019). Pytorch: An imperative style, high-performance deep learning library. Advances in neural information processing systems, 32.

Lehmann, D., & Skrandies, W. (1980). Reference-free identification of components of checkerboard-evoked multichannel potential fields. Electroencephalography and clinical neurophysiology, 48(6), 609–621.

