## Supplementary Information for "Spatiotemporal Decomposition of Whole-Brain Alpha Traveling Waves"

### Supplementary Note 1: Theoretical foundation of WOCCA

This section describes fundamental concepts and properties of the WOCCA framework. By proving our main theorem, the Traveling Energy Decomposition Theorem, we provide a rigorous explanation for WOCCA algorithm's capability to decompose traveling waves. First, we revisit the basic concepts.

**Definition 1**

The *conjugate contrast function* between two complex-valued $n$-vectors $u$ and $v$:

$$w(u,v) \coloneqq |u^*v|^2 - |u^*\bar{v}|^2$$

Where $\bar{u}$ denotes complex conjugate and $u^*$ denotes conjugate transpose of $u$. When $w(u,v) = 0$, $u$ and $v$ are *weakly orthogonal*.

When the conjugate contrast function is applied to a vector and itself (i.e. $v = u$), denote $\mathrm{TE}[u] \coloneqq w(u,u)$ as *traveling energy* of $u$.

A complex $n$-vector $u$ is a *pure traveling wave* if and only if $\mathrm{TE}[u] = |u|^4 > 0$; $u$ is a *pure standing wave* if and only if $\mathrm{TE}[u] = 0$.

Additionally, $u$ is a *unit wave* if and only if $|u| = 1$.

Below are a few properties that are easy to verify.

**Lemma 1**

Let $u$ and $v$ be complex $n$-vectors, we have:

a. $w(u,v) = w(v,u)$;
b. $w(u,\bar{v}) = -w(u,v)$;
c. For any complex number $c = ae^{i\theta}$ where $a$ and $\theta$ are real numbers, $w(u,cv) = a^2 w(u,v)$;
d. $-|u|^2|v|^2 \le w(u,v) \le |u|^2|v|^2$. Specifically, $0 \le w(u,u) \le |u|^4$.
e. Let $u = u_R + iu_I$, $u$ is a pure traveling wave if and only if $u_R$ and $u_I$ are orthogonal and $|u_R| = |u_I| > 0$.

**Proof:**

**a-d** are trivial and we only prove **e**. If $u$ is a pure traveling wave, $\mathrm{TE}[u] = |u|^4 > 0$, but:

$$\begin{aligned}\mathrm{TE}[u] &= |u^*u|^2 - |u^*\bar{u}|^2 \\ &= |u|^4 - |(u_R + iu_I)^*(u_R - iu_I)|^2 \\ &= |u|^4 - |(|u_R|^2 - |u_I|^2) + 2i \cdot u_R^T u_I|^2\end{aligned}$$

Real and imaginary part of the latter term need to both be 0, therefore satisfy:

$$\begin{cases}|u_R| = |u_I| \\ u_R^T u_I = 0\end{cases}$$

Meanwhile, since $|u| > 0$, we have $|u_R| = |u_I| > 0$, completing the forward proof.

Conversely, if $u_R^T u_I = 0$ and $|u_R| = |u_I| > 0$, backtrack the previous derivation we have $\mathrm{TE}[u] = |u|^4 > 0$.

■

It can be seen from **Lemma 1** that the conjugate contrast function is symmetric, invariant to constant phase shift, and that applying complex conjugate to one of the two vectors (equivalent to flipping traveling wave directionality) leads to negation of the function value, which is consistent with the properties of traveling waves. Meanwhile we notice that conjugate contrast function is not an inner product: it is only semi-positive definite and nonlinear in terms of scalar product. Therefore, a set of weakly orthogonal complex vectors don't form a set of linear bases. However, through a carefully designed transformation, we can interpret conjugate contrast function in a higher-dimensional space.

**Definition 2**

Consider function $f: \mathbb{C}^n \to \mathbb{R}^{n\times n}$:

$$f(u) \coloneqq \sqrt{2} Im[uu^*]$$

Function $f$ is the *skew-symmetric dimension-raising transform* of $u$, denoted in capital letter $U = f(u)$.

Define function $W$ as the sum of element-wise product of a square matrix:

$$W(U, V) \coloneqq \mathrm{tr}(U^T V)$$

And name $W$ as the *dimension-raised conjugate contrast function*.

As its name tells, the skew-symmetric dimension-raising transform converts a phasemap into a skew-symmetric matrix that is hopefully easier to deal with, and the element-wise dot product is the conjugate contrast function between a pair of phasemaps. We prove it in the following lemma.

**Lemma 2**

Let $u$ and $v$ be complex $n$-vectors, $\|U\|_F$ be Frobenius norm of $U$. We have:

a. For any complex number $c = ae^{i\theta}$ where $a$ and $\theta$ are real, $f(cu) = a^2 f(u)$;
b. $U$ is skew-symmetric, i.e. $U = -U^T$;
c. Rank of $U$ is either 0 or 2;
d. $w(u, v) = \sqrt{2}i \cdot v^* U v$;
e. $w(u, v) = W(U, V)$;
f. $\|U\|_F = \sqrt{W(U, U)} = \sqrt{\mathrm{TE}[u]}$.

**Proof:**

a. $(ae^{i\theta}u)(ae^{i\theta}u)^* = a^2 e^{i\theta} e^{-i\theta} \cdot uu^* = a^2 uu^*$.
b. Obviously $uu^*$ is Hermitian, therefore its imaginary part is skew-symmetric;
c. Each column of $Im[uu^*]$ is linear combination of the real and the imaginary part of $u$, therefore $\mathrm{Rank}[U] \leq 2$, but rank of a skew-symmetric matrix is always even, therefore it can only be 0 or 2.
d. Notice that:

$$Im[uu^*] = -\frac{1}{2} i \cdot \left[uu^* - \overline{(uu^*)}\right] = -\frac{1}{2} i \cdot (uu^* - \bar{u}\bar{u}^*)$$

And:

$$w(u, v) = w(v, u) = v^* u \cdot u^* v - v^* \bar{u} \cdot \bar{u}^* v = v^*(uu^* - \bar{u}\bar{u}^*)v$$

Substitute the middle term, we have:

$$w(u, v) = v^*\left(\sqrt{2}i \cdot U\right)v = \sqrt{2}i \cdot v^* U v$$

e. According to **d**:

$$\begin{aligned} w(u,v) &= \sqrt{2}i \cdot v^* U v \\ &= -\sqrt{2}\,\mathrm{Im}[v^* U v] \\ &= -\sqrt{2}\,\mathrm{Im}\left[\sum_{j,k} \bar{v}_j v_k U_{jk}\right] \\ &= -\sqrt{2}\,\mathrm{Im}\left[\mathrm{tr}\left(U^T(\bar{v}\bar{v}^*)\right)\right] \end{aligned}$$

Since $U$ is real, we only need to consider imaginary part of $\bar{v}\bar{v}^*$:

$$\begin{aligned} w(u,v) &= -\sqrt{2}\,\mathrm{tr}(U^T\,\mathrm{Im}[\bar{v}\bar{v}^*]) \\ &= \mathrm{tr}(U^T V) \\ &= W(U,V) \end{aligned}$$

f. Combining **e** and the definition of Frobenius norm and traveling energy, the proof is trivial.

■

The next theorem shows that not only every phasemap can be mapped to a low-rank skew-symmetric matrix, but also every low-rank skew-symmetric matrix corresponds to a unique pure traveling wave.

**Theorem 1 (Structure Theorem for Pure Traveling Waves)**

Define the set of low-rank skew-symmetric matrices $S_0$, and the subset $S$ with Frobenius norm equals 1 (plus the special case of zero matrix):

$$S_0 \coloneqq \{A | A \in \mathbb{R}^{n\times n}, A = -A^T, \mathrm{Rank}[A] \le 2\}$$
$$S \coloneqq \{A | A \in S_0, \|A\|_F = 1\} \cup \{0_{\{n\times n\}}\}$$

Multivalued mapping $g: S_0 \to \mathbb{C}^n$:

$$g(U) \coloneqq \begin{cases} 0_{\{n\}}, if\ \mathrm{Rank}[U] = 0 \\ Eigenvector\ of\ U\ with\ smallest\ imaginary\ egenvalue, if\ \mathrm{Rank}[U] = 2 \end{cases}$$

And the set $P_0$ of unit pure traveling waves and the zero vector:

$$P_0 \coloneqq \{a | a \in \mathbb{C}^n, |a| = \mathrm{TE}[u] = 1\} \cup \{0_{\{n\}}\}$$

$P_0$ is partitioned by the constant phase shift equivalent relation $a \sim e^{i\theta}a, \theta \in \mathbb{R}$, resulting in the quotient set $P$ of equivalent classes. We have:

a. $S_0$ is the image of $\mathbb{C}^n$ mapped by $f$;
b. If $\mathrm{Rank}[U] = 2$, $g(U)$ will be a unit pure traveling wave. Therefore $g$ becomes a function considering the above equivalent relation and $P$ is the image of $S_0$ mapped by $g$;
c. $S$ is the image of $P$ mapped by $f$. $f: P \to S$ and $g: S \to P$ are inverse to each other.

**Proof:**

a. **Lemma 2c** and **2d** have shown that the image of $\mathbb{C}^n$ mapped by $f$ is within $S_0$. We only need to prove that $f$ is surjective. The case of zero matrix is trivial. For rank-2 skew-symmetric matrix $U$, let $k > 0$ be absolute value of imaginary part of the only pair of non-zero imaginary eigenvalues and $z$ be the unit eigenvector corresponding to eigenvalue $ki$, matrix $U$ can be expressed as:

$$U = ki \cdot zz^* - ki \cdot \bar{z}\bar{z}^* = 2k \cdot \mathrm{Im}[\bar{z}\bar{z}^*]$$

Therefore, let $u = \sqrt{2k}\bar{z}$, we have:

$$f(u) = \sqrt{2}Im[uu^*] = 2k \cdot Im[\bar{z}\bar{z}^*] = U$$

b. Following the construction in **a**, we have $g(U) = \bar{z}$. Since $z$ and $\bar{z}$ are eigenvectors corresponding to two distinct eigenvalues, $z$ is orthogonal to $\bar{z}$ and $|z| = |\bar{z}| = 1$, and:

$$\mathrm{TE}[\bar{z}] = |\bar{z}^*\bar{z}|^2 - |z^*\bar{z}|^2 = |\bar{z}|^4 - 0 = 1$$

Therefore $g(U)$ is a unit pure traveling wave when $\mathrm{Rank}[U] = 2$. This shows that image of $S_0$ mapped by $g$ is within $P_0$. Since eigenvectors are compatible with constant phase shift equivalent relation, $g: S_0 \to P$ becomes a single-valued function.

We also need to prove that $g$ is surjective. The case of zero matrix is trivial. On the other hand, for pure traveling wave $u = u_R + iu_I$ in $P_0$, let:

$$U = f(u) = \sqrt{2}Im[uu^*]$$
$$= \sqrt{2}(u_I u_R^T - u_R u_I^T)$$

According to **Lemma 1e**:

$$Uu = \sqrt{2}[(u_I u_R^T u_R - u_R u_I^T u_R) + i(u_I u_R^T u_I - u_R u_I^T u_I)]$$
$$= \sqrt{2}\left[\left(u_I \cdot \frac{\sqrt{2}}{2} - u_R \cdot 0\right) + i\left(u_I \cdot 0 - u_R \cdot \frac{\sqrt{2}}{2}\right)\right]$$
$$= -iu_R + u_I$$
$$= -iu$$

This shows that $u$ is an eigenvector of $U$ with eigenvalue $-i$, while we already know $U \in S_0$ has only one pair of non-zero and conjugate imaginary eigenvalues, which are obviously $\pm i$. Therefore $u$ is in the same equivalent class as $g(U)$ and is in the image of $S_0$ mapped by $g$;

c. First, **Lemma 2a** shows that the value of $f$ is constant within a constant phase shift equivalent class of $P_0$, allowing a nature definition of $f$ on $P$. When $u$ is zero vector, $f(u)$ is zero matrix. When $u$ is a unit pure traveling wave, according to **Lemma 2f**, its Frobenius norm $\|f(u)\|_F = 1$. Therefore, the image of $P$ mapped by $f$ is within $S$.

We also need to prove that $f$ restricted to $P$ is still a surjection. The case of zero matrix is trivial. On the other hand, for non-zero matrix $U \in S$, following the construction in **a**, let:

$$u = \sqrt{2k}\bar{z}$$

And we have $U = f(u) = 2kf(\bar{z})$. We only need to prove $u$ is a unit pure traveling wave. We notice that $\bar{z}$ is a unit pure traveling wave, because:

$$\mathrm{TE}[\bar{z}] = |z|^4 - |z^*\bar{z}|^2 = |\bar{z}|^4$$

We also notice $\|U\|_F = 1$, while also:

$$\|U\|_F = 2k\|f(\bar{z})\|_F = 2k$$

Therefore $2k = 1$, which means $u = \bar{z}$ and is a unit pure traveling wave. In this way, the image of $P$ mapped by $f$ completely covers $S$.

Finally, the construction that we have used repeatedly has clearly shown that $f$ and $g$ are inverse to each other between sets $P$ and $S$.

■

The relationship between sets and mappings in **Theorem 1** is shown in the commutative diagram below, where $\pi$ is quotient map defined by constant phase shift equivalent relation and $i$ is identity embedding.

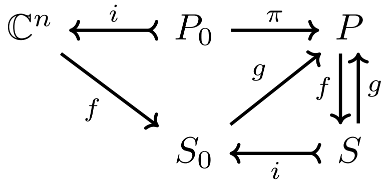


Regarding the conjugate contrast function, since $f$ and $g$ are inverse to each other between sets $P$ and $S$, according to **Lemma 2e**, function $w$ and $W$ are fully consistent between $P$ and $S$. It is worth noting that the results for $g$ function only applies to set $S$ rather than $S_0$, since $g$ maps all non-zero matrices to unit vectors. If Frobenius norm of the matrices doesn't equal 1, then (assuming $U$ and $V$ are non-zero matrices):

$$W(U,V) = \|U\|_F \|V\|_F \cdot W\left(\frac{U}{\|U\|_F}, \frac{V}{\|V\|_F}\right) = \|U\|_F \|V\|_F \cdot w\big(g(U), g(V)\big)$$

The relationship described above is shown in the commutative diagram below.

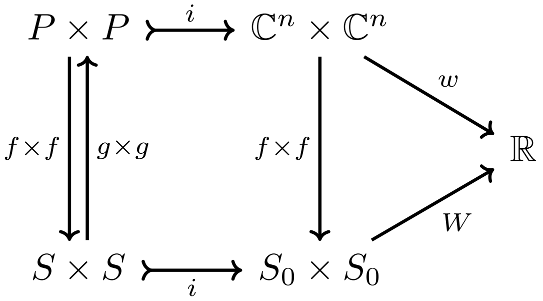


These results suggest that dimension raising to the space of skew-symmetric matrices enables more concise and computable expression of the conjugate contrast function. Indeed, the $W$ function is the standard element-wise inner product of matrices in space $\mathbb{R}^{n\times n}$. Consequently, two weakly orthogonal vectors in $\mathbb{C}^n$ become truly orthogonal after being transformed to $S_0$ with $W$ being the inner product. Furthermore, the norm derived from $W$ function is the Frobenius norm, therefore traveling energy is the square of norm in matrix space. **Theorem 1** shows that every phasemap can be transformed into either a pure traveling wave or zero vector (when the phasemap is a pure standing wave) by the composite mapping $f \circ g$ (referred to as the "purification mapping"). In the first case, the pure traveling wave is the "purified" traveling pattern in the original phasemap, and the square of Frobenius norm of the skew-symmetric matrix obtained from $f$ function is the traveling energy of the phasemap, standing for the traveling pattern's "intensity". Therefore, describing a phasemap through its low-rank skew-symmetric matrix via $f$ function is essentially using the pattern and magnitude (norm) of the matrix to represent the innate pure traveling pattern and its traveling energy of the phasemap, while eliminating all other factors and characteristics unrelated to traveling waves.

These insights inspired us to transfer the notion of standard orthogonal bases from Euclidean and unitary spaces to the context of traveling waves and conjugate contrast.

**Definition 3**

A set of complex $n$-vectors $\{u_1, u_2, \ldots, u_m\}$ is a set of *standard weakly orthogonal basis* if and only if:

a. For every $j = 1,2, \ldots, m$, $u_j$ is a unit pure traveling wave;

b. If $j \neq k$, $u_j$ and $u_k$ are weakly orthogonal, i.e. $w(u_j, u_k) = 0$.

Based on the definition, we prove the main theorem of the WOCCA framework to demonstrate that a standard weakly orthogonal basis specifically decomposes the traveling patterns in a set of phasemaps.

**Theorem 2 (Traveling Energy Decomposition Theorem)**

Given a set of complex $n$-vectors $D = \{v_1, v_2, \ldots, v_d\}$ and a standard weakly orthogonal basis $F = \{u_1, u_2, \ldots, u_m\}$ that also consists of $n$-vectors, define the *total traveling energy* of $D$:

$$\mathrm{TE}[D] \coloneqq \sum_j \mathrm{TE}[v_j]$$

And *traveling energy of* $D$ *explained by* $F$:

$$\mathrm{ETE}[D, F] \coloneqq \sum_{j,k} w(u_j, v_k)^2$$

The inequality below holds:

$$\mathrm{TE}[D] \geq \mathrm{ETE}[D, F]$$

**Proof:**

Convert all complex vectors into matrices in $S_0$ using $f$ function, all traveling energy and conjugate contrast function values remain unchanged, and total traveling energy becomes sum of squared norm of matrices:

$$\mathrm{TE}[D] = \sum_j \|V_j\|_F^2$$

Traveling energy explained by $F$ becomes the sum of squared inner product. We also denote the $W$ function in the form of inner product $\langle A, B \rangle \coloneqq \mathrm{tr}(A^T B)$, then:

$$\mathrm{ETE}[D, F] = \sum_{j,k} \langle U_j, V_k \rangle^2$$

Meanwhile, since $\|U_j\|_F^2 = \mathrm{TE}[u_j] = 1$ and $\langle U_{j_1}, U_{j_2} \rangle = w(u_{j_1}, u_{j_2}) = 0$ when $j_1 \neq j_2$, in matrix space $\{U_1, U_2, \ldots, U_m\}$ is a standard orthogonal basis of a subset. On this set of basis, the sum of squared norm (total variance) is obviously not less than sum of squared inner product.

■

Theorem 2 demonstrates that projecting a set of phasemaps on a set of weakly orthogonal basis via conjugate contrast function is essentially decompose traveling energy or "intensity" of traveling patterns across explained traveling energy by each component in the basis. To some extent, this is similar to a standard orthogonal basis decomposing total variance across explained variance by each component. However, contrary to the fact that total variance in a Euclidean space can always be full decomposed and the sum of explained variance equals total variance, a weakly orthogonal basis doesn't guarantee a full decomposition. To be precise, for a standard weakly orthogonal basis $F$ in **Theorem 2**, there doesn't always exist an expansion $\hat{F} = F \cup \{u_{m+1}, u_{m+2}, \dots, u_{m+p}\}$ that makes $\mathrm{TE}[D] = \mathrm{ETE}[D, \hat{F}]$, but the analogous expansion does exist for Euclidean and unitary spaces. This is because although the space of skew-symmetric matrices isn't completely spanned by $\{U_1, U_2, \dots, U_m\}$, it is possible that the orthogonal subspace of the spanned subspace doesn't contain any matrices with rank 2. In other words, rank-2 skew-symmetric matrices have been "depleted" by the current standard weakly orthogonal basis, thus the expansion cannot continue.

### Supplementary Note 2: Weak orthogonality and the condition of independent frame

It is important to explain why conjugate contrast function and its derived weakly orthogonal constraint, which plays a centric role in the WOCCA algorithm, defines independence between traveling wave patterns. In this section we prove an equivalent condition of weak orthogonality, the condition of independent frame. Below is a detailed formulation and proof.

**Theorem 3 (Condition of Independent Frame)**

Let $u$ and $v$ be complex $n$-vectors, and define the *set of static frames* $F[u]$ of $u$ as:

$$F[u] \coloneqq \{\mathrm{Re}[e^{i\theta}u] | \theta \in [0,2\pi)\}$$

Then $w(u,v) = 0$ if and only if the *condition of independent frame* holds:

$$\exists x \in F[u]\ \forall y \in F[v]\ x^T y = 0$$

**Proof:**

The set of static frames is linear combination of real and imaginary part of the complex vector. Let $u = u_R + iu_I$:

$$\mathrm{Re}[e^{i\theta}u] = u_R \cos\theta - u_I \sin\theta$$

First prove sufficiency. If the condition of independent frame holds, consider the cases of $\theta = 0$ and $\theta = \frac{3}{2}\pi$:

$$\exists x \in F[u] \begin{cases} x^T v_R = 0 \\ x^T v_I = 0 \end{cases}$$

By definition we can assume $x = \mathrm{Re}[e^{i\phi}u]$, then obviously $x^T v = 0$ and $x^T \bar{v} = 0$. Also let the imaginary part $p = \mathrm{Im}[e^{i\theta}u]$, we have:

$$\begin{aligned} w(u,v) &= w(e^{i\phi}u, v) \\ &= |(x+ip)^* v|^2 - |(x+ip)^* \bar{v}|^2 \\ &= |ip^T v|^2 - |ip^T \bar{v}|^2 \\ &= 0 \end{aligned}$$

Next prove necessity. Given $w(u,v) = 0$, let $z = e^{i\theta}u$, first we notice:

$$z_R^T v_R = u_R^T v_R \cos\theta - u_I^T v_R \sin\theta$$

It is a sinusoidal function of $\theta$, thus there must be some $\theta$ that satisfies $\mathrm{Re}[z]^T v_R = 0$. In this case:

$$\begin{aligned} w(z,v) &= |z_R^T v_R + z_I^T v_I + i(z_R^T v_I - z_I^T v_R)|^2 - |z_R^T v_R - z_I^T v_I - i(z_R^T v_I + z_I^T v_R)|^2 \\ &= (z_I^T v_I)^2 + (z_R^T v_I - z_I^T v_R)^2 - (z_I^T v_I)^2 - (z_R^T v_I + z_I^T v_R)^2 \\ &= 4 z_R^T v_I \cdot z_I^T v_R \end{aligned}$$

We know that $w(z,v) = w(u,v) = 0$, therefore $z_R^T v_I \cdot z_I^T v_R = 0$, which means either $z_R^T v_I = 0$ or $z_I^T v_R = 0$. The former implies that $z_R$ is orthogonal to both real and imaginary part of $v$. Since $z_R$ is a static frame of $u$, the condition of independent frame holds between $u$ and $v$. The latter implies that $v_R$ is orthogonal to both real and imaginary part of $z$. The sets of static frames are identical for $z$ and $u$, therefore every static frame of $u$ is orthogonal to $v_R$. We can find another phase $\phi$ at which $\mathrm{Re}[e^{i\phi}u]$ and $v_I$ are orthogonal (because $\mathrm{Re}[e^{i\phi}u]^T v_I$ is also a sinusoidal function of $\phi$), in this case $\mathrm{Re}[e^{i\phi}u]$ is orthogonal to both the real part and the imaginary part of $v$. This proves the condition of independent frame between $u$ and $v$.

■

The condition of independent frame means that for a pair of weakly orthogonal phasemaps, any of them contains at least one “key frame” orthogonal to all static frames of the other phasemap. This provides a figurative and intuitive guarantee that the spatial modes of traveling wave propagation represented by the pair of phasemaps are truly unique to each other.

### Supplementary Note 3: Construction and properties of the WOCCA algorithm

This section describes the process of WOCCA algorithm in detail, as well as a few important properties that supports its rationality. Projection on standard weakly orthogonal basis is essentially decomposing the traveling energy (i.e. intensity of traveling patterns) of the phasemap data. Based on the objective of explaining more traveling energy in the leading components and utilizing the symmetry of elements in skew-symmetric matrices to reduce the dimensionality of matrix space, we define the WOCCA algorithm as below.

#### Algorithm 1 (WOCCA)

Given a set of complex $n$-vectors $D = \{v_1, v_2, \dots, v_d\}$:

i. Transform vectors in $D$ into skew-symmetric matrices $\{V_1, V_2, \dots, V_d\}$ using function $f$, and express them using their upper triangle elements in the form of $\mathbb{R}^{\frac{n(n-1)}{2}}$;

ii. Compute the covariance matrix $B \in \mathbb{R}^{\frac{n(n-1)}{2} \times \frac{n(n-1)}{2}}$ of these upper triangle element vectors. Specifically:

$$B = \sum_j \mathrm{up.tri.}\left[f(v_j)\right] \mathrm{up.tri.}\left[f(v_j)\right]^T$$

Where $\mathrm{up.tri.}[A]$ stands for the upper triangle element vector form of $A$;

iii. Initialize set of complex $n$-vectors $F = \emptyset$, and counting variable $j = 1$;

iv. Find a non-zero complex $n$-vector $u_j$ that satisfies the weak orthogonality constraint:

$$\forall u_k \in F \; w(u_j, u_k) = 0$$

And maximize the objective function:

$$Obj := \frac{1}{|u_j|^4} \cdot \mathrm{up.tri.}\left[f(u_j)\right]^T B \, \mathrm{up.tri.}\left[f(u_j)\right]$$

The objective function is actually $\frac{1}{4}\mathrm{ETE}\left[D, \{u_j\}\right]$ where the factor $\frac{1}{4}$ comes from choosing only the upper triangle elements;

v. If the optimal objective is greater than 0, add $\frac{u_j}{|u_j|}$ to set $F$ and $j \leftarrow j + 1$, then go back to **step iv**;

vi. If the optimal objective is 0 or no valid $u_j$ exist, terminate the algorithm and return set $F$.

The result of the algorithm has properties described in the following theorem.

#### Theorem 4

Let set $F$ be the decomposition result from WOCCA algorithm applied to a certain set of complex $n$-vectors. We have:

a. $F$ is a set of standard weakly orthogonal basis;

b. For vectors $u_j$ and $u_k$ corresponding to indices $j$ and $k$, if $j < k$ then:

$$\mathrm{ETE}\left[D, \{u_j\}\right] \geq \mathrm{ETE}[D, \{u_k\}]$$

c. If the algorithm didn't terminate due to optimal objective being 0, then $F$ contains at least $n-1$ vectors.

**Proof:**

a. First, vectors in $F$ are weakly orthogonal to each other and $|u_j| = 1$. We only need to prove that $u_j$ is a pure traveling wave for all $j$s. We prove it by contradiction. Assume $u_j$ is not a pure traveling wave, since the objective is not 0, we know that $f(u_j)$ is not zero matrix. We construct a pure traveling wave $\widehat{u_j} = g\left(f(u_j)\right) \in P$ and prove $\frac{1}{\|f(u_j)\|_F} f(u_j) \in S$:

$$\frac{1}{\|f(u_j)\|_F} f(u_j) = f\left(g\left(\frac{1}{\|f(u_j)\|_F} f(u_j)\right)\right)$$
$$= f\left(g\left(f(u_j)\right)\right)$$
$$= f(\widehat{u_j})$$

Since $\|f(u_j)\|_F = \sqrt{\mathrm{TE}[u_j]} < 1$, we can substitute $u_j$ with $\widehat{u_j}$ and the corresponding skew-symmetric matrix is multiplied by a factor greater than 1, making the non-zero objective function larger. This implies that $u_j$ is actually not optimal and contradicts the definition of the algorithm.

b. Trivial.

c. We only need to prove that if $F$ contains only $n-2$ vectors or less, a new pure traveling wave vector that orthogonal to all existing vectors can be added to it. Assume $|F| = k$, since:

$$\mathrm{Rank}[u_{R1}, u_{R2}, \dots, u_{Rk}] \le k \le n-2$$

We can find a two-dimensional subspace orthogonal to all these real-valued vectors, and construct a standard orthogonal basis $\{p, q\}$ of it. We only need to take $u_{k+1} = \frac{\sqrt{2}}{2}(p + iq)$, by **Lemma 1e** $u_{k+1}$ is a pure traveling wave. Also when $j \le k$, the real and imaginary part of $u_{k+1}$ are all orthogonal to $u_{Rj}$, therefore the set of static frames of $u_{k+1}$ is orthogonal to $u_{Rj}$. By **Theorem 3**, $u_{k+1}$ is weakly orthogonal to $u_j$, hence $u_{k+1}$ is indeed a valid expansion.

It is worth noting that, strengthening the proposition to $n$ vectors does not hold. When $n = 2$ the dimension of the space of all skew-symmetric matrices is only 1, therefore the number of vectors in $F$ cannot exceed 1.

■

The above properties guarantee that WOCCA can separate independent traveling wave components from phasemap data, and greedily explain the total traveling energy in data by the leading components, which is consistent with PCA. Meanwhile, since the dimensionality of phasemap data (i.e. number of channels) is high, **Theorem 4c** makes sure that a relatively large number of leading components can be obtained before the WOCCA algorithm stops.

### Supplementary Note 4: CPCA and standing waves

This section proves a lemma about pure standing waves based on the WOCCA framework, and explain a disadvantageous property of CPCA when dealing with traveling wave data. That is, CPCA prefers standing waves when processing pairs of traveling waves with opposite directionality.

**Lemma 3**

Let $u$ and $v$ be complex $n$-vectors, we have:

a. If $u$ is a pure standing wave, there exists a $\theta \in \mathbb{R}$ satisfying $e^{i\theta}u \in \mathbb{R}^n$;
b. If $u$ is a pure standing wave, $w(u,v) = 0$ holds for any $v$;
c. $u$ is a pure standing wave if and only if $f(u) = 0_{\{n\times n\}}$.

**Proof:**

a. $\mathrm{TE}[u] = |u|^4 - |\bar{u}^*u|^2$, therefore $|u|^4 = |\bar{u}^*u|^2$. The inequality:
$$|\bar{u}^*u| \le |u|^2$$
Equality holds if and only $\bar{u}$ and $u$ are colinear, i.e. there exists a $\theta \in \mathbb{R}$ that satisfies $\bar{u} = e^{2i\theta}u$. Therefore:
$$e^{i\theta}u = e^{-i\theta}\bar{u} = \overline{e^{i\theta}u}$$
Which means $e^{i\theta}u$ is real-valued.
b. $\mathrm{ETE}[\{u\},\{v\}] = w(u,v)^2$, but $\mathrm{ETE}[\{u\},\{v\}] \le \mathrm{TE}[u] = 0$.
c. If $u$ is a pure traveling wave, take $\theta$ as defined in **a**, then:
$$f(u) = f\left(e^{i\theta}u\right) = 0$$
Conversely, if $U = f(u) = 0_{\{n\times n\}}$, we have:
$$\mathrm{TE}[u] = w(u,u) = W(U,U) = 0$$

■

The disadvantageous property of CPCA when decomposing paired traveling wave data is described as below.

**Theorem 5**

Let $D = \{v_1, v_2, \dots, v_d\}$ be a set of complex $n$-vectors. Denote:
$$\widehat{D} = \{v_1, \overline{v_1}, v_2, \overline{v_2} \dots, v_d, \overline{v_d}\}$$
Then under the optimization objective of CPCA applied to set $\widehat{D}$:
$$Obj \coloneqq \frac{1}{|u|^2}\sum_{v\in\widehat{D}} |u^*v|^2$$
The objective can always reach its maximum at a pure standing wave.

**Proof:**

The optimization objective can be written as:
$$Obj = \frac{1}{|u|^2}\sum_{v\in D}(|u^*v|^2 + |\bar{u}^*v|^2)$$
It is obviously invariant to scaling of $i$, therefore we can assume $|u| = 1$.

First, if $u$ is a unit pure standing wave, from **Lemma 3a** there exists a $\theta \in \mathbb{R}$ satisfying $e^{i\theta}u \in \mathbb{R}^n$. The objective function is also invariant to constant phase shift, therefore we can assume $u$ is real-valued. In this case, let $\widetilde{D} = \{v_{R1}, v_{I1}, v_{R2}, v_{I2}, \dots, v_{Rd}, v_{Id}\}$:

$$\begin{aligned} Obj &= 2\sum_{v\in D}|u^*v|^2 \\ &= 2\sum_{v\in D}(|u^Tv_R|^2+|u^Tv_I|^2) \\ &= 2\sum_{x\in\widetilde{D}}|u^Tx|^2 \end{aligned}$$

The objective function of WOCCA is twice the PCA objective function applied to set $\widetilde{D}$;

On the other hand, if $u$ is not a pure standing wave, then:

$$\begin{aligned} Obj &= \sum_{v\in D}(|u_R^Tv_R+u_I^Tv_I+i(u_R^Tv_I-u_I^Tv_R)|^2+|u_R^Tv_R-u_I^Tv_I+i(u_R^Tv_I+u_I^Tv_R)|^2) \\ &= 2\sum_{v\in D}[(u_R^Tv_R)^2+(u_I^Tv_I)^2+(u_R^Tv_I)^2+(u_I^Tv_R)^2] \\ &= 2\sum_{x\in\widetilde{D}}[(u_R^Tx)^2+(u_I^Tx)^2] \end{aligned}$$

In this case, denote $y$ as the first principal component of real-valued PCA applied to $\tilde{d}$. We can write $u_R$ and $u_I$ as linear combination of $y$ and other components orthogonal to $y$: $u_R=\alpha_Ry+\beta_Rz_R$, $u_I=\alpha_Iy+\beta_Iz_I$, where $z_R,z_I\in\mathbb{R}^n$ and $\alpha_R^2+\beta_R^2+\alpha_I^2+\beta_I^2=1$, $|y|=|z_R|=|z_I|=1$, $y^Tz_R=y^Tz_I=0$. Then:

$$Obj=2\sum_{x\in\widetilde{D}}[(\alpha_R^2+\alpha_I^2)(y^Tx)^2+\beta_R^2(z_R^Tx)^2+\beta_I^2(z_I^Tx)^2]$$

By the fundamental properties of PCA (where $z$ can be $z_R$, $z_I$ or any other real vector satisfying $|z|=1$):

$$\sum_{x\in\widetilde{D}}(y^Tx)^2\ge\sum_{x\in\widetilde{D}}(z^Tx)^2$$

Therefore:

$$Obj\le 2\sum_{x\in\widetilde{D}}|y^Tx|^2$$

But previously we have proven that equality can holds when $u$ is a unit pure standing wave.

■

Theorem 5 demonstrates that CPCA cannot distinguish between standing waves and pairs of conjugate traveling waves, hence only possible to identify traveling patterns with highly asymmetric dimensionality. Although a pure traveling wave and its complex conjugate are actually orthogonal ($\mathrm{TE}[u]=|u|^4$ is equivalent to $|\bar{u}^*v|=0$), CPCA, built for standard orthogonal basis, is surprisingly incapable of the orthogonal conjugate pair that exists in the data. On contrary, WOCCA is designed specifically for traveling waves, and describe each pair of opposite traveling directions using one weak orthogonal component, therefore has an obvious advantage over CPCA.

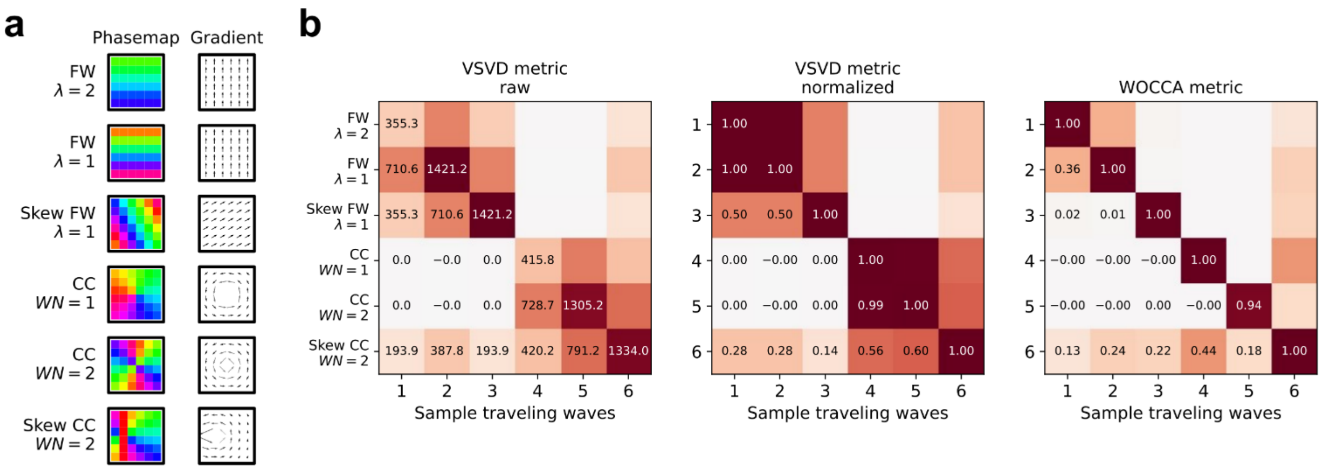


**Supplementary Figure 1: VSVD cannot account for both wavelength and pattern similarity at the same time**

VSVD measures similarity between two traveling wave patterns using inner product between phase gradient fields. Magnitude of vectors in a phase gradient field measures the magnitude of local phase gradients, the reciprocal of which is equivalent to wavelength of a traveling wave. Therefore, magnitude of inner product reflects both pattern similarity between two phase gradient fields and their innate characteristics such as wavelengths (for traveling waves with clear propagation direction) or winding numbers (for rotational waves).

a. Typical phasemaps and corresponding phase gradient fields synthesized on a 6x6 grid, including forward longitudinal waves with long wavelength, short wavelength, and skewed short wavelength, as well as counterclockwise rotational waves with winding number 1, winding number 2, and skewed winding number 2. The purpose of constructing skewed traveling waves is to test whether a traveling wave metric can separate the contribution of wavelength from spatial pattern similarity.
b. Left: simply using inner product of phase gradient fields can identify different wavelengths, but comparing traveling wave no. 1 with 3 or no. 4 with 6, we notice that traveling waves with short wavelength and low pattern similarity cannot be separated from those with long wavelength and high pattern similarity. Middle: the normalized inner product metric that we actually used in practice can identify different level of pattern similarity, but by comparing traveling wave no. 1 with 2 or no. 4 with 5, we notice that it loses the ability to identify wavelengths due to normalization. Right: conjugate contrast metric of WOCCA can clearly distinguish all traveling waves in this example.

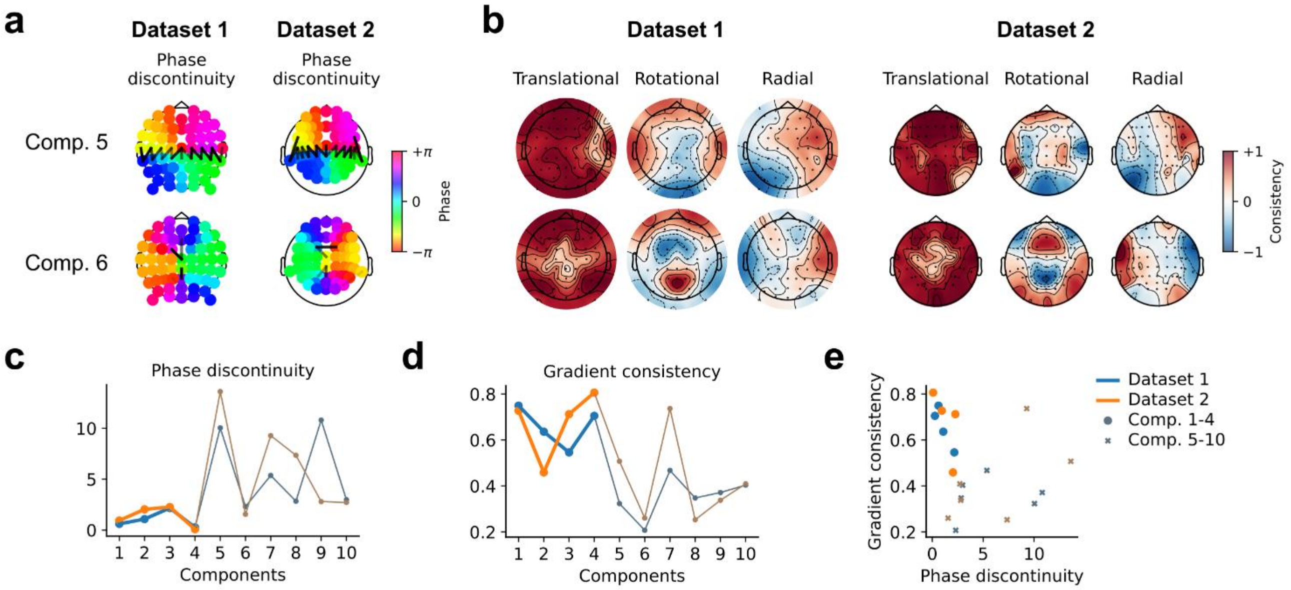


**Supplementary Figure 2: Choosing the number of WOCs**

We define typical traveling wave structures as a combination of "translational type" with locally coherent propagation direction and "singularity type" that is rotationally symmetric with respect to a center point (including rotations, sources, drains, etc.). These structures exhibit spatial continuity of both phase and phase gradient, except for locations close to the singularities.

a. Local continuity of phase is defined by whether phase differences between neighboring electrodes (determined by triangulation of electrode layout) are greater than $90°$. In Dataset 1 and 2, spatial pattern of phase of Component 5 is not coherent between anterior and posterior regions.
b. Local continuity of phase gradient field is defined by local structure's goodness of fit of translational type and singularity type standard structures, where the singularity type consists of rotational singularity and radial singularity (including sources and drains). At the horizontal boundary between anterior and posterior brain, Component 6's goodness of fit is low for both translational and singularity types.
c. We define the sum of squares of the part of phase difference greater than $90°$ as the quantitative index of phase discontinuity. The first four components all exhibit very low discontinuity, while the following components, especially component 5, do not.
d. For each electrode, we define the maximum of its neighbor's goodness of fit of translational, rotational, and radial structures as the quantitative continuity of local phase gradient. For the whole phase gradient field, we take the global minimum as the quantitative index of phase gradient continuity. The first four components all exhibit very high continuity, while the following components, especially component 5, do not.
e. We use phase discontinuity and phase gradient continuity as the main indices for measuring consistency of traveling waves' spatial structures. The first four components of both datasets are grouped at the top left corner of the plane, suggesting high consistency.

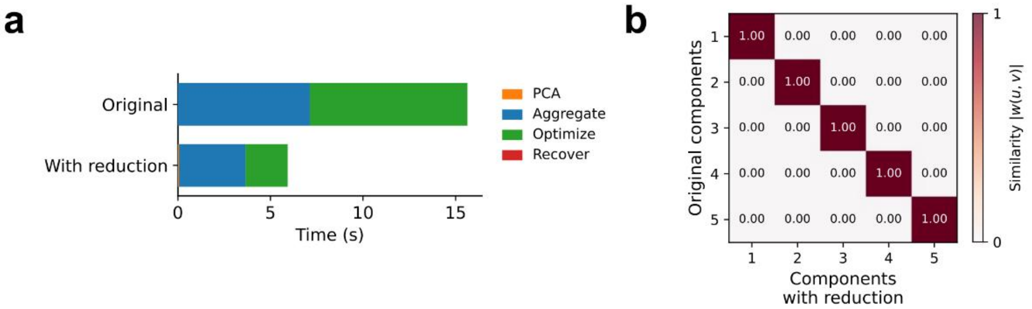


**Supplementary Figure 3: Efficiency and precision of preliminary dimensionality reduction**

We synthesized longitudinal and rotational traveling waves on a larger 8x8 grid (5,000 samples for each direction of each type, 20,000 in total), and compared results between a direct WOCCA and applying a 30-dimensional PCA dimensionality reduction before WOCCA.

a. Although pre-reduction introduced additional steps of PCA in the beginning and reprojection in the end, pre-reduced WOCCA takes much less time compared to direct WOCCA. The efficiency in both aggregating quadratic form matrices and nonlinear optimization improves greatly.

b. The first five WOCs yielded by pre-reduced WOCCA and direct WOCCA match well and are practically identical (measured using absolute value of conjugate contrast).

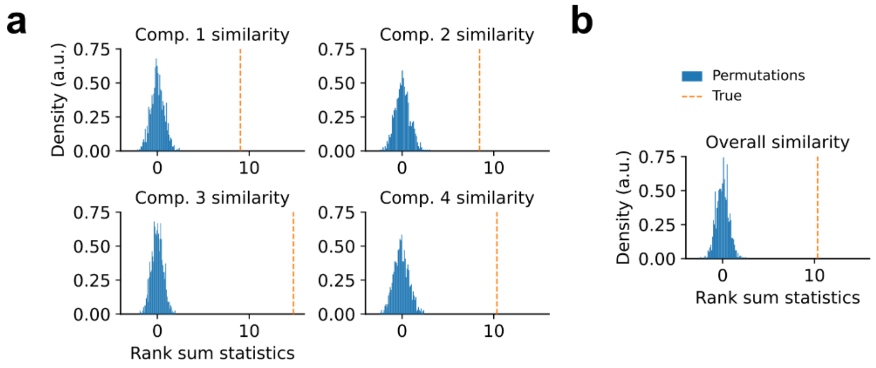


**Supplementary Figure 4: Non-parametric test of within subject stability of WOCs**

a. Randomly shuffle subject identities and session orders in Dataset 2, and apply WOCCA to each session of each subject for 1,000 times. Wilcoxon rank-sum statistics are calculated between within-subject and between-subject similarity. The empirical value is greater than all 1,000 shuffled samples in the null distribution.

b. Conducting the same test on the average similarity of the four components.

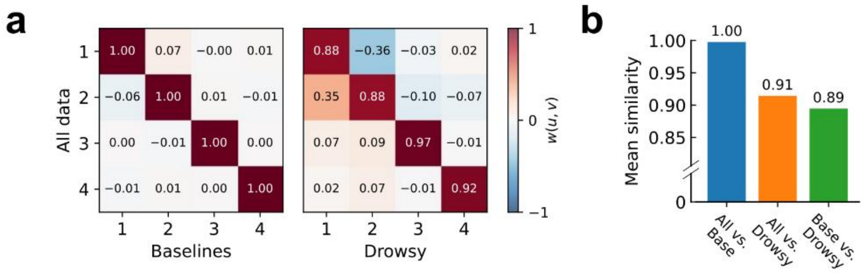


**Supplementary Figure 5: Comparing unsupervised WOCCA results between different sedation levels**

a. Similarity of first four WOCs comparing all data and baseline stage data (left), and all data and "drowsy" state data (right). WOCs from all data is similar to baseline stage data, while WOCs from both all data and baseline stage data are significantly different from "drowsy" state data (the latter comparison see **Fig. 5b** in main text)

b. Average of diagonal elements in all three similarity matrices.

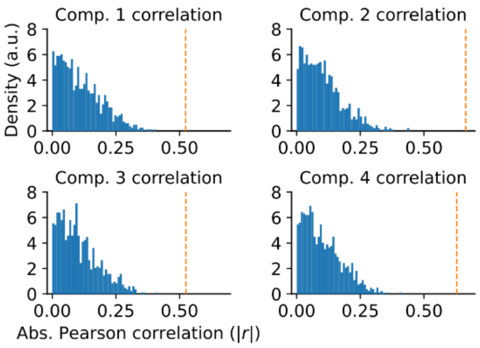


**Supplementary Figure 6: Non-parametric test of correlation between blood propofol concentration and traveling waves**

Randomly shuffle blood propofol concentrations of all subjects at all stages, and apply the supervised WOCCA weighted on propofol levels for 1,000 times. Absolute value of Pearson correlation coefficients between traveling energy of each component and propofol levels were calculated. The empirical value is greater than all 1,000 shuffled samples in the null distribution.

| Traveling energy between EC and EO, Dataset 1 | | | | |
|---|---|---|---|---|
| **Components** | Rot. | Long. | Horiz. | Lat. |
| **P-value** | $0.092$ | $6.0 \times 10^{-24}$ | $7.5 \times 10^{-16}$ | $0.063$ |
| **Cohen's d** | $0.119$ | $0.809$ | $-0.615$ | $0.131$ |
| **Result** | N.S. | *** EC>EO | *** EC<EO | N.S. |
| **Traveling energy between EC and EO, Dataset 2** | | | | |
| **Components** | Rot. | Long. | Horiz. | Lat. |
| **P-value** | $4.9 \times 10^{-41}$ | $2.1 \times 10^{-69}$ | $0.055$ | $0.12$ |
| **Partial $\eta^2$** | $0.453$ | $0.646$ | $0.0123$ | $0.00796$ |
| **Result** | *** EC>EO | *** EC>EO | N.S. | N.S. |

**Supplementary Table 1: Comparison of traveling energy between EC and EO resting states**

| Direction asymmetry between EC and EO, Dataset 1 | | | | |
|---|---|---|---|---|
| **Components** | Rot. (CC) | Long. (FW) | Horiz. (RW) | Lat. (FS) |
| **P-value** | $0.66$ | $0.0024$ | $2.4 \times 10^{-6}$ | $4.2 \times 10^{-9}$ |
| **Cohen's d** | $0.0313$ | $-0.215$ | $0.341$ | $-0.431$ |
| **Result** | N.S. | ** EC<EO | *** EC>EO | *** EC<EO |
| **Direction asymmetry between EC and EO, Dataset 2** | | | | |
| **Components** | Rot. (CC) | Long. (FW) | Horiz. (RW) | Lat. (FS) |
| **P-value** | $0.022$ | $6.1 \times 10^{-6}$ | $1.7 \times 10^{-5}$ | $4.3 \times 10^{-16}$ |
| **Partial $\eta^2$** | $0.0175$ | $0.0662$ | $0.0600$ | $0.199$ |
| **Result** | *EC<EO | *** EC<EO | *** EC>EO | *** EC<EO |

**Supplementary Table 2: Comparison of directionality asymmetry between EC and EO resting states**

| Direction asymmetry non-zero, Dataset 1 | | | | |
|---|---|---|---|---|
| **Components** | EC, Rot. | EC, Long | EC, Horiz | EC, Lat |
| **P-value** | $0.52$ | $1.7 \times 10^{-11}$ | $0.0084$ | $7.0 \times 10^{-15}$ |
| **Cohen's d** | $-0.0448$ | $-0.501$ | $0.187$ | $-0.591$ |
| **Result** | N.S. | *** FW<BW | ** RW>LW | *** FS<BS |
| **Components** | EO, Rot. | EO, Long. | EO, Horiz. | EO, Lat. |
| **P-value** | $0.076$ | $1.3 \times 10^{-14}$ | $0.0046$ | $0.0044$ |
| **Cohen's d** | $-0.125$ | $-0.584$ | $-0.201$ | $-0.202$ |
| **Result** | N.S. | *** FW<BW | ** RW<LW | ** FS<BS |
| **Direction asymmetry non-zero, Dataset 2** | | | | |
| **Components** | EC, Rot. | EC, Long. | EC, Horiz. | EC, Lat. |
| **P-value** | $0.16$ | $0.0038$ | $0.032$ | $2.8 \times 10^{-8}$ |
| **Cohen's d** | $-0.183$ | $-0.389$ | $0.283$ | $-0.826$ |
| **Result** | N.S. | ** FW<BW | * RW>LW | *** FS<BS |
| **Components** | EO, Rot. | EO, Long. | EO, Horiz. | EO, Lat. |
| **P-value** | $0.28$ | $0.0026$ | $0.83$ | $8.2 \times 10^{-4}$ |
| **Cohen's d** | $-0.140$ | $-0.406$ | $-0.0276$ | $-0.455$ |
| **Result** | N.S. | ** FW<BW | N.S. | *** FS<BS |

**Supplementary Table 3: Significance of directional asymmetry at resting state**

For rotational waves, CC: counterclockwise, CW: clockwise. For longitudinal waves, FW: forward, BW: backward. For horizontal waves, RW: rightward, LW: leftward. For lateral waves, FS: forward on sides, BS: backward on sides.

One-sample t test was used to test the significance of directional asymmetry, comparing it to 0 (i.e. the two directions of the traveling wave pattern have identical intensity).

$*: P < 0.05$; $**: P < 0.01$; $***: P < 0.001$.
